# Levels of additive genetic variation vary substantially between species

**DOI:** 10.64898/2026.01.22.701036

**Authors:** Lillith Zijmers, Katie Abson, Jarrod Hadfield, Adam Eyre-Walker

## Abstract

A population’s ability to adapt is determined by its levels of additive genetic variance (V_A_), and while it is agreed that most organisms have genetic variation for most traits, the extent to which it varies between species is poorly characterised. Here we investigate this question by compiling 3209 and 1852 estimates of heritability and evolvability (the additive genetic variance divided by the square of the mean) estimates respectively, for a variety of traits, from 220 and 172 multicellular eukaryotic species. Using phylogenetic generalised linear mixed models, we find substantial and highly significant interspecific variation in evolvability. Much of the variation is explained by phylogenetic relatedness, with plants in our data having substantially higher evolvability than animals. While heritability also varies between species, the differences are more subtle, and plants are not exceptional. We investigate whether the variation in evolvability and heritability between species is due to variation in the mutation rate, effective population size, genome size, ploidy and recombination rate, but find little evidence of any factor being important. However, the confidence intervals are large suggesting that we have little power to detect any associations between these factors and our estimates of V_A_.

## Introduction

A population’s ability to respond to selection depends on its levels of quantitative genetic variation or more specifically the additive genetic variation (V_A_) underlying key traits. It is well known that most traits have at least some underlying additive genetic variation (Hill *et al*., 2008; Falconer and Mackay, 2009) with substantial variation between published estimates. However not much is known about whether levels vary systematically between species, or about the factors influencing the observed variation in levels of quantitative genetic variation. Understanding this is key to understanding the genetic architecture underlying complex traits and their evolution.

Estimates of V_A_ are not generally comparable between traits and species because they are measured on different scales. Consequently, V_A_ is typically standardised by dividing the estimate by the phenotypic variance (V_P_), yielding the narrow-sense heritability (h^2^) (Falconer and Mackay, 2009). Historically, the primary focus on heritability arose from the use of the breeder’s equation in predicting the response to selection (Falconer and Mackay, 2009; Hansen and Pélabon, 2021). However, relying solely on heritability has some disadvantages for understanding the forces affecting quantitative genetic variance (QGV) because the additive genetic variance appears in both the numerator and the denominator: h^2^= V_A_ / (V_A_ + V_NA_ +V_E_), where V_NA_ is the non-additive genetic variance and V_E_ is the environmental variance. Furthermore, some of the other variance components in the denominator are likely to be correlated with V_A_ (Hansen *et al*., 2011). Therefore, an increase in V_A_ may not lead to a proportional increase in h^2^. In light of these issues and to better understand the additive genetic variance and thereby the adaptive potential of a population, the evolvability (I_A_), quantified as V_A_ divided by the square of the mean, was introduced by Houle (1992); note that in some analyses the square root of the evolvability is taken to yield the coefficient of additive genetic variance (CV_A_).

Although DNA sequence diversity is known to vary substantially between species (Leffler *et al*., 2012), there is little evidence of variation in V_A_ (Mittell *et al*., 2015). A recent large-scale analysis across birds and mammals found significant variation among species in heritability but not CV_A_ (Young and Postma, 2023) a pattern the authors suggested as being due to variation between species in the level of non-additive genetic or environmental variance, rather than V_A_. Previous comparative studies examining relationships between measures of additive genetic (I_A_ and h^2^) and microsatellite diversity, census population size, or ecological specialization also found no clear associations which would be indicative of variation in V_A_ (Mittell *et al*., 2015; Wood *et al*., 2016; Martinossi-Allibert *et al*., 2017). However, there are some cases in which V_A_ has been shown to vary between species for particular traits - for example, desiccation and cold resistance, but not wing size, across *Drosophila* species (Kellermann *et al*., 2009). Here, we compile estimates of I_A_ and h^2^ from a much broader range of multicellular eukaryotes than has been analysed before and investigate whether there is significant variation between species.

We also investigate whether I_A_ and h^2^ are correlated with various factors expected to influence the level of additive genetic variance. Most models for the maintenance of quantitative genetic variation predict that V_A_ should depend on the mutational input V_M_ (Walsh and Lynch, 2018). Although there is no evidence that estimates of V_M_ differ significantly between species, such estimates are relatively few and subject to substantial measurement error (Conradsen *et al*., 2022). Therefore, in this study we examine whether V_A_ is correlated with the nucleotide mutation rate and with the product of the mutation rate and proteome size, which represents the expected rate of mutations that might affect quantitative genetic variation. We use these as proxies for mutational input given the limited number of V_M_ estimates. We also investigate the effects of genome size, ploidy levels and the number of genes, as these factors may influence the effective mutational target size – the total number of sites at which mutations can alter quantitative traits - beyond what is captured by proteome size alone and thus the mutational input contributing to additive genetic variation (Walsh and Lynch, 2018; Besnard *et al*., 2020). Levels of V_A_ are also expected to depend on the effective population size when the trait is neutral (Lynch and Hill, 1986; Abson *et al*., 2025) or selection on the trait is weak (Walsh and Lynch, 2018; Bürger *et al*., 1989; Keightley and Hill, 1990). Wood et al. (2016) found no significant correlation between heritability (h^2^) and the census population size, however h^2^ is a poor comparative measure of V_A_ and census and effective population sizes are thought to be only moderately correlated (Buffalo, 2021; James and Eyre-Walker, 2020). We test whether V_A_ is correlated to long-term estimates of *N*_*e*_, estimated by dividing the nucleotide diversity by a direct measure of the mutation rate. Additionally, we might expect V_A_ to be influenced by the rate of recombination because recombination mediates the buildup and breakdown of linkage disequilibrium underlying the Bulmer effect (Bulmer, 1980), mitigates the reduction of variation caused by Hill-Robertson interference among linked selected sites (Hill and Robertson, 1966; McVean and Charlesworth, 2000), and limits the formation of pseudo-overdominance due to linked recessive alleles (Ohta, 1971; Charlesworth and Willis, 2009). We explore this by correlating estimates of V_A_ against the recombination rate per nucleotide, along with whether the species is obligately sexual, or engages in asexual reproduction for some parts of its lifecycle. We also examine whether additional biological variables—such as body mass, body length, longevity, and generation time—are correlated with our measures of additive genetic variance.

## Methodology

### Literature search and QGV compilation

Heritability and evolvability estimates were compiled from the peer-reviewed literature indexed in the Web of Science for the period between 1992 and 2022. Searches were conducted using “Heritability” or “Evolvability” as topic keywords, with the aim of capturing estimates from a diverse range of species and from natural populations. Therefore, journals focusing on domesticated, crop species or human populations were excluded. The search was further filtered by only selecting journals with over 200 papers meeting the search criteria and then scanning the first 50 articles by titles when sorted by relevance. This resulted in the choice of four journals: ‘Journal of Evolutionary Biology’, ‘Evolution’, ‘Heredity’ and ‘Proceedings of the Royal Society B’. When multiple estimates were available for the same trait within a population, we selected a single value based on a standardised evaluation of reliability. Preference was given to estimates derived from larger sample sizes and model structures minimising potential biases. When estimates were otherwise comparable in these respects, we applied a hierarchy based on the relatedness structure used to estimate additive genetic variation. Animal models were preferred, as they utilise information from all known relationships and are generally the least biased and most precise (Akesson *et al*., 2008). When animal model estimates were unavailable, we selected those from parent-offspring regressions, which are less affected by dominance or common-environmental effects than sibling-based designs (Hill, 2009; Lynch and Walsh, 1998).

The resulting dataset was then merged with the datasets of Mittell *et al*. (2015) and Young and Postma (2023). All recorded estimates of evolvability and heritability were numerically checked where possible and recalculated if sufficient raw data were available. Each estimate was also evaluated for suitability in terms of the trait scale and transformation, as evolvability is only meaningful for traits measured on ratio or log-interval scales, where proportional differences are interpretable (Houle, 1992; Hansen *et al*., 2011). Heritability estimates were likewise checked for compatibility with the underlying trait scale, as h^2^ is interpretable for interval and ratio-scale traits, but not for ordinal or nominal traits unless modelled and interpreted appropriately (Falconer and Mackey, 1996; Lynch and Walsh, 1998). If an evolvability estimate was not given, this was calculated if the necessary data were provided. We removed estimates where artificial selection had been performed on the focal trait, along with estimates from inbred lines, since inbreeding inflates V_A_. However, estimates were retained from species that are naturally clonal or selfing. We log_10_ transformed estimates of I_A_ because the estimates are not normally distributed. Consequently, 146 negative and zero evolvability values were removed, as these are undefined under log transformation.

For cases in which both I_A_ and h^2^ have been estimated we can calculate the residual variance, the sum of the non-additive and environmental variances, which we normalised by dividing it by the square of the mean: I_R_=I_A_(1-h^2^)/h^2^ (Hansen *et al*., 2011).

### Interspecific Variation Models

To partition the variation in recorded estimates of V_A_ a phylogenetic generalised linear mixed model (PGLMM) was fitted using Markov chain Monte Carlo (MCMC) implemented in MCMCglmm (Hadfield, 2010) in R (R Core Team, 2023). Weakly informative priors were used throughout. Random-effect variances were modelled using scaled F_1,1_ priors (scale √1000), while the residual variance followed an inverse-gamma prior with shape and scale set to 0.001. Fixed effects were assigned normal priors with a mean 0 and variance 10^8^. The MCMC chains were run for 780,000 iterations, with a burn-in of 180,000 and thinning every 250 iterations. Model convergence was assessed through visual inspection of chain mixing. Additionally, we used posterior predictive checks to visually assess model fit. We included a number of fixed and random effects in our model which aim to capture the likely major sources of variation in V_A_ (Table 1). Critical to our analysis are two species-level random effects that capture the variation in V_A_ among species, one associated with phylogenetic inertia (“Phylogeny” in Table 1) and one which is independent of this (“Non-Phylogenetic” in Table 1). The phylogenetic term takes into account that closely related species may covary in their levels of V_A_; the tree was estimated using Time Tree (Kumar *et al*., 2022).

**Table 1.** Summary of the fixed and random effects included in all MCMCglmm models, with the levels of the categorical fixed effects and the rationale for inclusion. For the random effects the number of levels differ between the models for evolvability and heritability and these are separated by | (I_A_|h^2^).

|  | Levels | Explanation |
| --- | --- | --- |
| Fixed effects |  |  |
| Method | 7 | Estimates of $V_A$ can differ across estimation methods because study designs and statistical models can vary in how they partition genetic and environmental sources of variation, and because they rely on different type of relatives (Young and Postma, 2023; Walsh and Lynch, 1998) |
| Trait Type | 4 | Estimates of $V_A$ depend on trait type (e.g. morphological, life-history) (Young and Postma, 2023; Mittell *et al.*, 2015, Hansen *et al.*, 2011) |
| Dimension | 6 | Assigned based on mathematical properties of the focal trait. Evolvabilities depend on trait dimension (Hansen *et al.*, 2011) |
| Environment | 3 | Estimates vary between Lab and Wild populations (Hansen and Pélabon, 2021) |
| Covariates |  |  |
| Number of fixed effects | Continuous | Included to reflect potential systematic variation in QGV due to model complexity |
| Number of random effects | Continuous |  |
| Random effects |  |  |
| Publication ID | 269\|415 | Indexes the publications contributing data. Each estimate is assigned a study ID corresponding to the publication it originated from, accounting for non-independence among estimates reported within the same paper |
| Trait | 652\|1036 | Categorical variable representing conceptually similar traits. Trait names vary across studies - and some measures captured slightly different aspects of the same underlying phenotype - we standardised trait labels across the dataset. Groups similar traits to account for inherent similarities accounting for inherent similarities |
| **Phylogeny** | 172\|220 | Accounts for variation in $V_A$ that is due to shared evolutionary history among species |
| **Non-Phylogenetic** | 172\|220 | Captures species-specific variation in $V_A$ that is not explained by phylogenetic relatedness |

The model was run for evolvability and heritability separately. Estimates of the predicted log evolvability and heritability for each species were calculated by extracting the intercept, phylogeny and species estimates for the *i*th species at the *j*th sampling of the chain

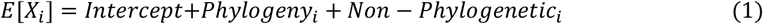

Where X is either log (I_A_) or h^2^. These were then averaged across samples of the chain to yield the posterior mean evolvability and heritability for a linear morphological trait estimated with an animal model in a wild population (these are the reference categories in our analysis), along with the 95% credible intervals.

To test whether there is significant variation between specific phylogenetic groups represented in our dataset, we averaged the species estimates within each taxonomic subgroup for each sampling of the MCMC. We then calculated pairwise differences between these group means (e.g. plants vs. animals, arthropods vs. mammals etc.). From these contrasts we derived the posterior distribution of the difference in means, quantifying the variation among the phylogenetic groups sampled in our data.

To further investigate whether phylogenetic groups differ in their average evolvability or heritability we estimated the evolvabilities and heritabilities for the internal nodes (*n*) of our phylogeny, providing posterior estimates for ancestral nodes.

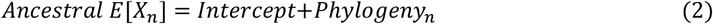

These node-level estimates were then used to calculate pairwise differences among major clades, following the same procedure as above. Finally, we extended our original interspecific variation models by including taxonomic groups as fixed effects, allowing us to formally test whether differences in log (I_A_) or h^2^ between groups deviate more than expected under the null model of Brownian evolution (the phylogenetic term) with added independent and identically distributed species effects (the non-phylogenetic term).

To quantify the amount of variation between our species in terms of evolvability and heritability we took the estimated variance associated with the phylogenetic (V_phylogenetic_) and non-phylogenetic (V_non-phylogenetic_) species terms and then assuming the terms are normally distributed we derived the quartiles of the distribution of the combined distribution. We quantify the variation in terms of the ratio of the upper and lower quartiles. This allows us to state for example, that the top 25% of species have an evolvability that is at least x-fold greater than the bottom 25% of species. The 95% credible intervals on these ratios were derived from the samples of the chain.

### Correlation analysis

We investigated whether our estimates of I_A_ and h^2^ could be predicted by a number of factors: the point mutation rate, the point mutation rate multiplied by the number of protein coding sites in the genome, genome size, number of genes, ploidy levels, the effective population size, the rate of recombination per nucleotide, the mode of reproduction, body mass, body length, longevity, and generation time. Body length was defined as the longest linear dimension; we restricted this analysis to animals since body length is hard to assess in plants because of the root system. Data on the predictors were taken from various databases, including Amniote (Myhrvold *et al*., 2015), Pantheria (Jones *et al*., 2009), Animal Diversity Web (Myers *et al*., 2025) as well as some meta-analytic studies such as Wang and Obbard (2023), Lynch et al. (2023), Stapley *et al*. (2017) and the broader literature (see Table S1 for details). All predictors were log_10_ transformed prior to analysis.

To assess the predictive power of each of our predictors, we used the same fixed and random effect structure, prior and iteration settings as for the interspecific variation models. Since, we did not have estimates of each predictor for every species, we ran the analysis for each predictor separately with heritability and evolvability respectively as the response variable. The amount of between-species variance explained by each predictor was quantified using a signed R^2^ (Mittell *et al*., 2015).

Where V(x) is the variance in the focal predictor and β is the regression coefficient. The product β^2^V(x) represents the between-species variation in the respective measure of V_A_ that is attributable to variation in the predictor, with the sign of R^2^ taken from the sign of β. The denominator quantifies the total between-species variation, including phylogenetic and species-level components.

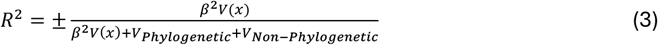

pMCMC values were used to assess the degree to which coefficients overlapped zero (Baayen et al., 2008) and for multi-level factors we used the logic of the Wald test to perform an omnibus test of the factor’s overall effect. The Wald statistic was calculated from the posterior mean vector and covariance matrix (Mittell *et al*., 2015).

### Evolvability versus Heritability

To investigate the relationship between heritability and evolvability we fitted a bivariate PGLMM with the phylogenetic and non-phylogenetic terms as random effects (Table 1) in which h^2^ and I_A_ were modelled jointly. The phylogenetic effects, non-phylogenetic effects and residuals were each allowed to covary across response variables and priors were placed on the resulting covariance matrices such that the marginal prior for the variances was identical to that in the univariate models. All MCMC settings were identical to those of the univariate models.

There is an expectation for V_A_ to be positively correlated with the non-additive genetic variance and the environmental variance (Hansen *et al*., 2011). The sum of these two components is the residual variance (V_R_), which we scale by dividing by the square of the mean to yield I_R_. To test whether I_A_ and h^2^ are correlated to I_R_ we ran bivariate PGLMMs under the same conditions as the h^2^ and I_A_ model.

## Results

To assess whether additive genetic variance varies systematically between species we compiled a dataset of evolvability (I_A_), and heritability (h^2^) estimates from the literature. Our dataset comprises 1,852 evolvability estimates from 172 species and 3,209 heritability estimates from 220 species, spanning a broad range of multicellular eukaryotes (Figure 1 and 2). We have estimates from roughly equal numbers of birds, mammals, arthropods and plants, with a small number of estimates from fish, arachnids, amphibians and reptiles. Many of our estimates were for linear morphological traits estimated using an animal model (493 I_A_ and 572 h^2^ estimates). For all variables aside from heritability we perform the analysis of the logarithm of the value, however, we refer to the variable by its name – e.g. all analyses are performed on log_10_(I_A_), but we refer to I_A_ and evolvability throughout the text.

**Figure 1.**
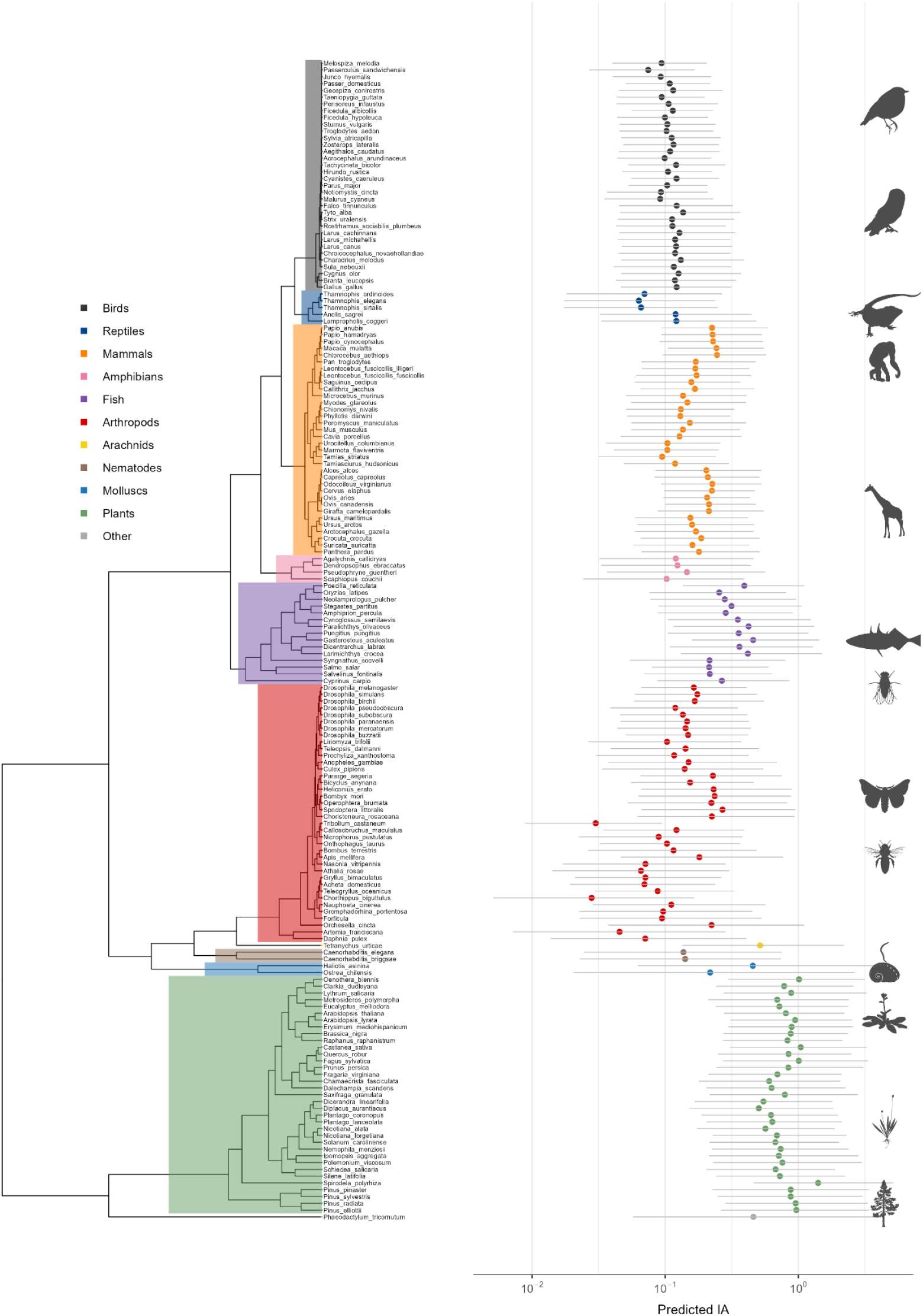
The predicted evolvability from our model for a linear, morphological trait estimated with an animal model from a wild population for each species, plotted across the phylogeny with their 95% credible intervals.

**Figure 2.**
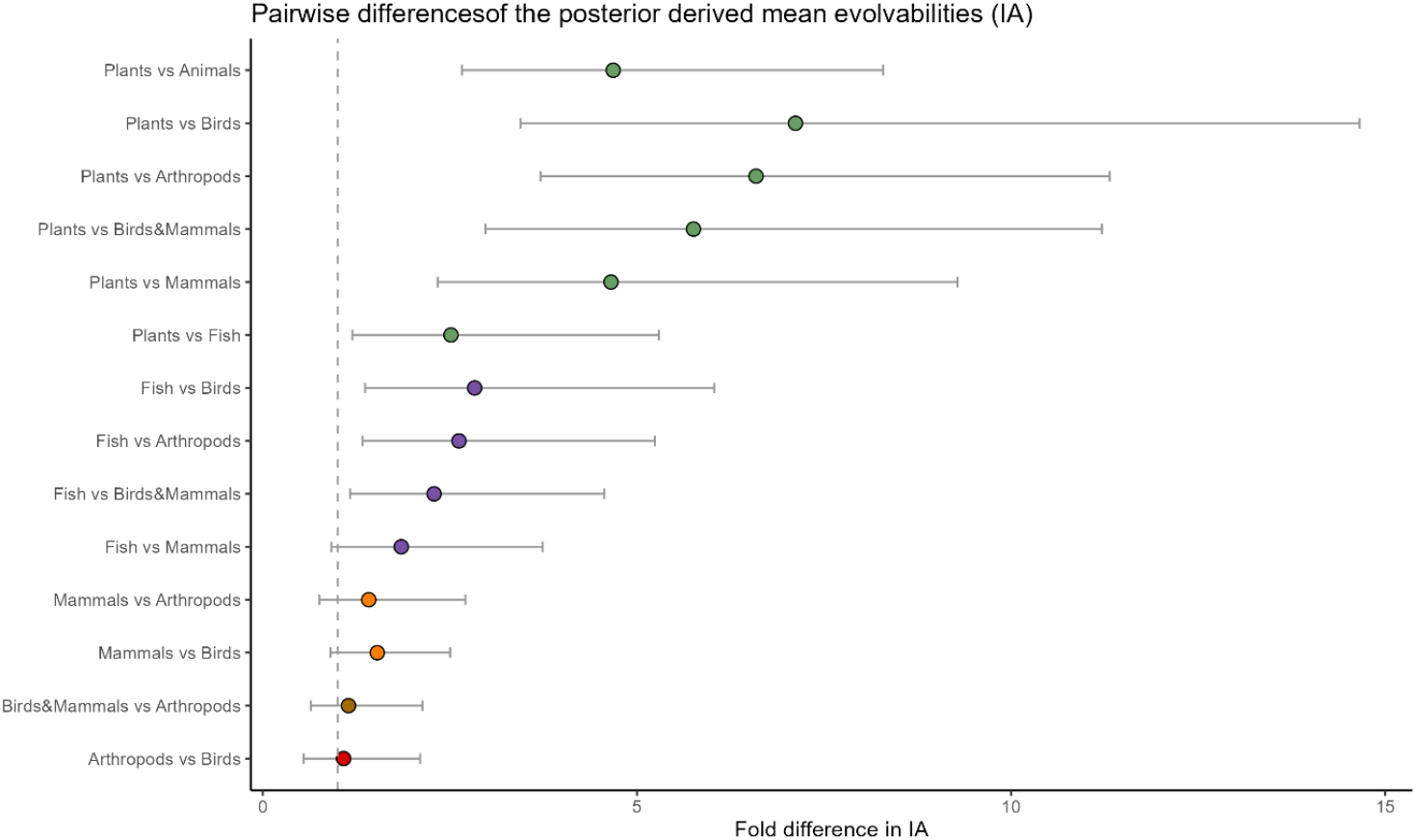
The difference in mean evolvability between groups of animals and plants. Each point represents the posterior mean difference between group means, back transformed to the original data scale. Horizontal bars indicate the 95% credible intervals. The dashed line at 1 indicates no difference between groups. Differences greater than 1 indicate that the first node (first-mentioned group in the y-axis) has a x-fold higher predicted evolvability than the second node. All comparisons are plotted so that the means are positive.

### Evolvability (I_A_)

To investigate whether I_A_ varies between species we fit a phylogenetic GLMM with two species terms – one capturing phylogenetic differences and another capturing species-specific deviations independent of phylogeny. The fit of the model to the data was adequate, although the model predicted a deficit of small values and an excess of large values compared to that observed (Figure S1). We find that I_A_ varies significantly between species (Figure 1), with 39% [95% credible intervals: 16% to 58%] of the total variance in the random effects (including the residual variance) in the model attributable to interspecific differences (Table 2). The magnitude of this variation is substantial: the top 25% of species exhibit 9.3-fold [3.5, 23] greater evolvability than the bottom 25%. Most of the interspecific variation reflects phylogenetic structure, with phylogeny accounting for 38% [15%, 58%] (Table 2) of the random effect variance.

**Table 2.** The proportion of the random effects variance explained by the phylogenetic and non-phylogenetic species variances for evolvability. Also given is the ratio of the upper and lower quartiles of the combined phylogenetic and non-phylogenetic species effects on the data scale where 1 indicates no difference; estimates are given to two significant figures unless the value is <0.01, in which case it is listed as 0

| Species | #species | #estimates | Phylo | Non-phylo | Phylo+ non-phylo | Ratio of quartiles |
| --- | --- | --- | --- | --- | --- | --- |
| All | 172 | 1852 | 38%<br>[15, 58] | 1.0%<br>[0, 3.9] | 39%<br>[17, 58] | 9.3<br>[3.5, 23] |
| Animalia | 135 | 1395 | 22%<br>[4.7, 40] | 1.4%<br>[0, 5.4] | 24%<br>[6.6, 40] | 5.0<br>[2.5, 12] |
| Arthropods | 38 | 460 | 17%<br>[0, 40] | 6%<br>[0, 21] | 23%<br>[3.3, 48] | 5.8<br>[1.9, 20] |
| Birds | 34 | 360 | 4.7%<br>[0, 17] | 3.7%<br>[0, 13] | 8.4%<br>[0, 24] | 2.5<br>[1.1, 6.3] |
| Birds & Mammals | 68 | 698 | 12%<br>[0, 35] | 1.8%<br>[0, 7.0] | 14%<br>[0.01, 36] | 2.9<br>[1.2, 8.9] |
| Fish | 15 | 133 | 22%<br>[0, 68] | 22%<br>[0, 60] | 44%<br>[1.4, 83] | 7.2<br>[1.3, 63] |
| Mammals | 34 | 338 | 19%<br>[0, 43] | 3.3%<br>[0, 13] | 22%<br>[1.6, 46] | 3.1<br>[1.5, 6.8] |
| Plants | 35 | 425 | 13%<br>[0, 40] | 8.1%<br>[0, 22] | 21%<br>[0.16, 46] | 2.9<br>[1.2, 7.6] |

A substantial part of this phylogenetic effect appears to be a difference between plants and animals, with plants having 4.7-fold [2.7, 8.3] higher I_A_ than animals. Plants also have significantly higher evolvability than each of the individual animal subgroups (Figure 2). This difference between plants and animals, and individual animal groups is also evident in the estimated ancestral values for each group; the ancestor to our sample of plants is estimated to have had 4.3-fold [1.1, 18] higher evolvability than the ancestor of the sample of animals (Figure S2). However, the difference between plants and animals is consistent with the phylogenetic model. When group identity (plant vs. animal) is included as a fixed effect alongside the phylogenetic and non-phylogenetic species effects in the model, the fixed effect coefficient was not significantly different from zero (mean effect = 0.73 [-1.46, 2.52]; Table S3); i.e. plants are no more divergent from animals than we might expect given the time that has elapsed since the groups diverged and the rate at which evolvability changes.

However, the variation in evolvability between species is not simply due to the difference between plants and animals. If we restrict our interspecific variation model to animals only, we still find significant variation between species, with 24% [7%, 40%] of the variance attributable to species differences (Table 2). We also find significant variation in I_A_ between species within plants, mammals, arthropods, and fish individually (Table 2). In each of these cases, however, the phylogenetic and non-phylogenetic effects are not individually significant. This indicates that while there is a significant species effect, the data lack sufficient power to partition the species effect between the phylogenetic and non-phylogenetic effects.

Given the complexity of the full interspecific variation model, which includes many fixed and random effects, and an assumption that log (I_A_) is linearly related to the effects in our model, we performed a simplified analysis using only linear morphological traits estimated with an animal model. This results in 493 estimates from 63 species. Results were qualitatively unchanged: interspecific variation accounted for 58% [32%, 81%] of the random effect variance, and the phylogeny accounted for 56% [28%, 80%]. Moreover, if we simply plot the species’ mean I_A_ values for this data the phylogenetic effect remains evident (Figure S3).

Although, our principal focus is one whether evolvability varies between species, a number of other fixed effects are significant in our model. We observe substantial differences in evolvability among different trait types (Omnibus test, P(>x^2^) < 0.001), with behavioural traits exhibiting substantially greater I_A_ than other traits. We also find a significant effect of dimension (P(>x^2^) < 0.001; Table S4), where traits that scale quadratically (e.g. area) and cubically (e.g. mass) show 1.7 [0.83, 3.5] and 3.5 [2.2, 5.5] times greater evolvability than those that scale linearly (Figure S4). Counts exhibit 3.4 [1.9 - 5.9] times greater evolvability than linear traits. The method of estimation also has a significant effect (P(>x^2^) = 0.012; Table S4), with full-sib designs yielding higher estimates that are 4.0 [1.6, 9.8] times greater on average than animal models, and midparent-offspring regression yielding the lowest values that are 0.40 [0.17, 0.92] of the values from the animal model (Figure S4). Estimates of I_A_ also depend on the environment (P(>x^2^) = 0.003) in which the estimation was made. However, no individual level differed significantly from the reference level, and direct comparisons between laboratory and wild estimates were not significant (Table S4; Figure S4). This indicated that the overall significance arises from moderate differences across multiple environment categories rather than a single strong pairwise contrast.

Although these fixed effects account for some of the observed variation in I_A,_, they primarily serve to control for differences in methodology and trait choice across publications, and do not explain the interspecific variation in I_A_ that we detect. To investigate what might be causing this variation we conducted phylogenetic generalised linear mixed models (PGLMMs) including a suite of predictors: nucleotide mutation rate, estimated total point mutation rate of the proteome, genome size, number of genes, ploidy level, recombination rate per site, mode of reproduction, and estimates of effective population size. In addition, we assessed associations between evolvability and life-history traits such as body mass, body length, longevity, and generation time.

Among all variables tested, only body length (which we have only assessed for animals) exhibited a statistically significant association with evolvability after applying a Bonferroni correction for multiple testing. Body length accounted for approximately 47% [6.6%, 95%] of the between-species variance in evolvability, and the model predicts that doubling body size increases I_A_ by 1.3-fold [1.1, 1.5]. No other predictors were significant predictors of evolvability after correction for multiple tests; however, the credible intervals are broad indicating we have little power to detect associations (Table 3).

**Table 3.** The regression of evolvability on various variables. Given is the number of estimates and the number of species from which those estimates come from for each analysis, along with the posterior means with their CIs and p_MCMC_ (twice the posterior probability that the effect lies on the opposite site of zero from its estimated direction) as well as the signed R^2^ for all I_A_ indicating the proportion of the between-species variance explained. For the categorical predictors the p-value of the omnibus test are denoted as P(>X^2^). For the categorical predictors, the posterior means are the deviation from the reference levels (Diploid and Sexual, respectively). An * in the p_MCMC_ or P(>χ^2^) column indicates a significant effect after Bonferroni correction.

| Predictor of $I_A$ | N estimates | N species | Post. mean | $p_{MCMC}$ | $R^2$ |
| --- | --- | --- | --- | --- | --- |
| **Mutation Rate** | 514 | 43 | -0.385<br>[-0.828, 0.040] | 0.076 | -0.108<br>[-0.340, 0.024] |
| **Protein Coding Mutation Rate** | 413 | 31 | -0.432<br>[-0.881, 0.028] | 0.060 | -0.225<br>[-0.770, 0.024] |
| **Genome Size** | 918 | 64 | -0.109<br>[-0.542, 0.254] | 0.582 | -0.027<br>[-0.186, 0.069] |
| **Number of Genes** | 1223 | 104 | -0.043<br>[-0.987, 0.099] | 0.923 | -0.007<br>[-0.186, 0.069] |
| **Rate of Recombination** | 618 | 33 | 0.184<br>[-0.276, 0.663] | 0.433 | 0.054<br>[-0.092, 0.267] |
| **Effective Population Size** | 403 | 32 | 0.213<br>[-0.312, 0.728] | 0.416 | 0.038<br>[-0.058, 0.190] |
| **Generation Time** | 443 | 37 | -0.098<br>[-0.410, 0.186] | 0.496 | -0.113<br>[-0.679, 0.168] |
| **Longevity** | 1318 | 104 | -0.063<br>[-0.231, 0.119] | 0.483 | -0.019<br>[-0.106, 0.034] |
| **Body Length** | 868 | 79 | 0.370<br>[0.149, 0.588] | 0.003* | 0.471<br>[0.066, 0.951] |
| **Body Mass** | 751 | 71 | 0.107<br>[-0.020, 0.210] | 0.083 | 0.365<br>[-0.008, 0.920] |
| **Categorical Predictors** | **N estimates** | **N species** | **Post. mean** | **P(>$\chi^2$)** | **R<sup>2</sup>** |
| **Haplodiploid** | 1645 | 127 | 0.074<br>[-0.444, 0.593] | 0.67 | 0.029<br>[0.001, 0.105] |
| **Polyploid** |  |  | -0.035<br>[-0.431, 0.381] |  |  |
| **Mixed Reproduction** | 1784 | 159 | 0.208<br>[-0.107, 0.533] | 0.13 | 0.077<br>[0.002, 0.240] |
| **Asexual Reproduction** |  |  | 0.146<br>[-1.553, 1.806] |  |  |

### Heritability

When we run our model on heritability, we find that the model fits the data reasonably well except that there is a large excess of h^2^ estimates at zero compared to what the model predicts (Figure S5). This excess of zero estimates is almost certainly methodological, arising because many of the analyses return zero estimates of V_A_ when relatives exhibit negatively correlated trait values.

As with evolvability, we find evidence of significant variation between species – the proportion of the random effects variance explained by between species differences is 24% [4.9%, 52%]. However, this variation cannot be clearly attributed to either phylogenetic or non-phylogenetic components when considered separately (Table 4). The lack of a clear phylogenetic signal is evident when the predicted h^2^ from our model for each species are plotted on a phylogeny (Figure 3). We also find significant variation in h^2^ within plants and animals respectively, however the smaller taxonomic subgroups as mammals and fish only harbour marginally significant interspecific variation, while the remaining groups tested do not show significant differences (Table 4). Overall, we find that h^2^ varies quite substantially between species; the ratio of the upper and lower quartiles is 1.8-fold [1.3, 2.6].

**Table 4.** The proportion of the random effects variance explained by the phylogenetic and non-phylogenetic species variances for heritability. Also given is the ratio of the upper and lower quartiles of the combined phylogenetic and non-phylogenetic species effects on the data scale where 1 indicates no difference; estimates are given to two significant figures unless the value is <0.01, in which case it is listed as 0

| Species | #species | #estimates | Phylo | Non-phylo | Phylo+ non-phylo | Ratio of quartiles |
| --- | --- | --- | --- | --- | --- | --- |
| All | 222 | 3209 | 28%<br>[0.35, 57] | 4.4%<br>[0, 12] | 32%<br>[11, 60] | 1.8<br>[1.3, 2.6] |
| Animalia | 173 | 2497 | 12%<br>[0.67, 26] | 1.9%<br>[0, 6.3] | 14%<br>[2.1, 27] | 1.4<br>[1.2, 1.7] |
| Arthropods | 49 | 884 | 3.1<br>[0, 13] | 2.5%<br>[0, 9.7] | 5.6%<br>[0, 17] | 1.2<br>[1.0, 1.5] |
| Birds | 44 | 606 | 3.5<br>[0, 14] | 2.1%<br>[0, 7.8] | 5.6%<br>[0, 18] | 1.2<br>[1.0, 1.5] |
| Birds & Mammals | 82 | 1118 | 6.7%<br>[0, 25] | 1.4%<br>[0, 5.4] | 8.2%<br>[0, 26] | 1.2<br>[1.0, 1.6] |
| Fish | 17 | 225 | 21%<br>[0, 62] | 15%<br>[0, 44] | 36%<br>[0.01, 70] | 1.9<br>[1.1, 3.4] |
| Mammals | 38 | 512 | 19%<br>[0, 50] | 6.6%<br>[0, 25] | 25%<br>[0.15, 55] | 1.5<br>[1.1, 2.1] |
| Plants | 41 | 599 | 29%<br>[0, 67] | 10%<br>[0, 33] | 40%<br>[9.3, 71] | 2.1<br>[1.3, 3.6] |

**Figure 3.**
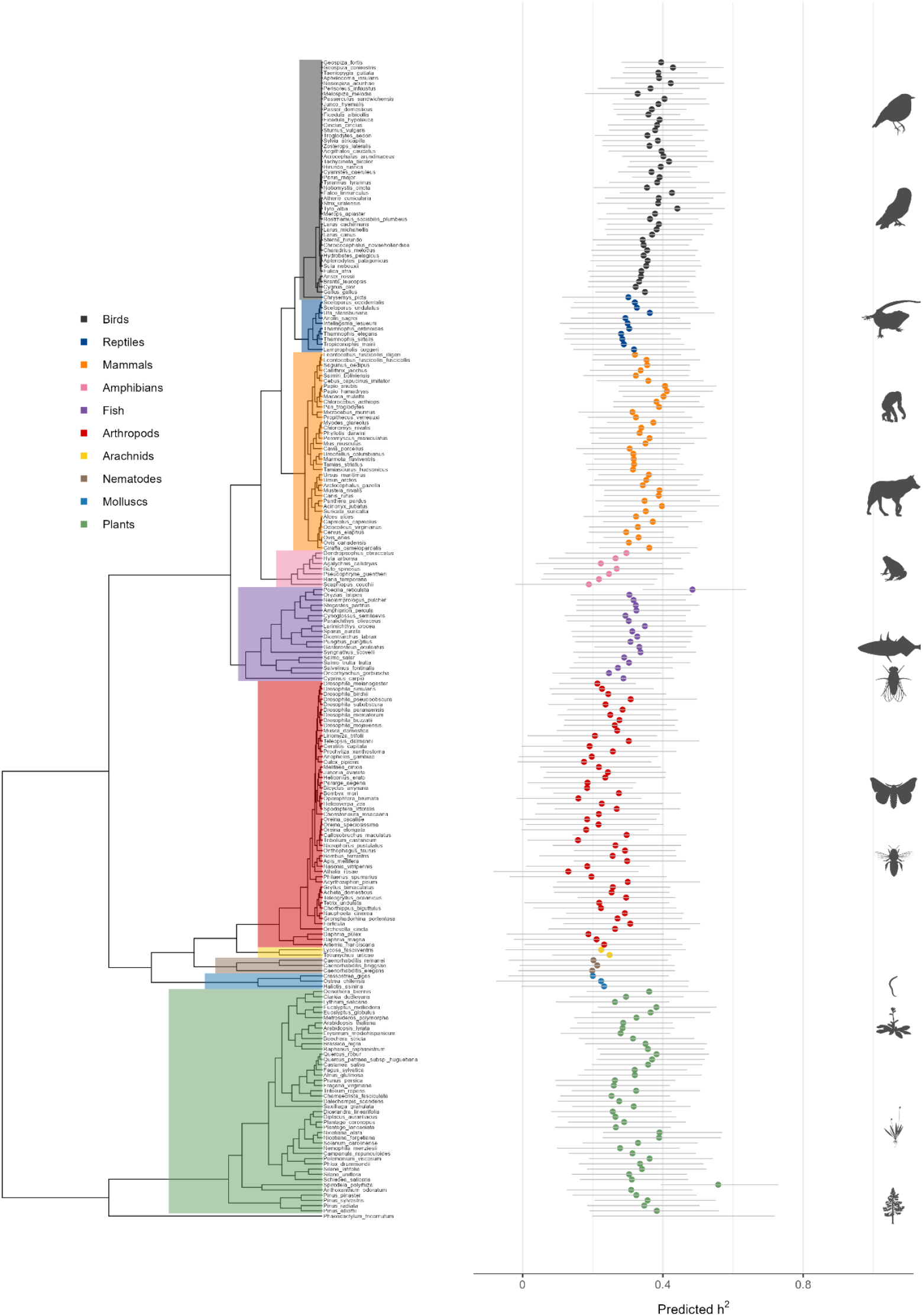
The predicted heritability from our model for a linear, morphological trait estimated with an animal model from a wild population for each species, plotted across the phylogeny with their 95% credible intervals.

All the fixed effects (Table 1) in our interspecific variation model (i.e. the model without any predictors such as the nucleotide mutation rate) are significant, with the exception of the number of fixed effects (p_MCMC_ = 0.118; Table S5). Consistent with previous research, h^2^ varies significantly between trait types (Omnibus test, P(>x^2^) < 0.001) with morphological traits having the highest h^2^ and fitness traits the lowest (the average difference in heritability between morphological and fitness traits is 0.15 [0.013, 0.28] (Table S5; Figure S6). We also find that the method of estimation has a significant effect (p_MCMC_ < 0.001, Table S5; Figure S6) with full-sib designs yielding estimates of heritability that are 0.21 [0.11, 0.30] higher than animal models, which yield similar estimates to other methods. We also find a weak but significant effect of dimension (P(>x^2^) = 0.012; Table S5; Figure S6) and the environment (P(>x^2^) = 0.002, Table S5; Figure S6). The effect of the environment seems to largely reflect higher heritability estimates from laboratory studies compared to those from natural populations (p_MCMC_ =0.003, Table S5; Figure S6).

Although we observe significant variation in narrow-sense heritability (h^2^) across species, most predictors tested do not exhibit statistically significant associations after Bonferroni correction (Table 5). The exception is ploidy level (P(>x^2^) = 0.004), with polyploids showing lower heritability than diploids.

**Table 5.**
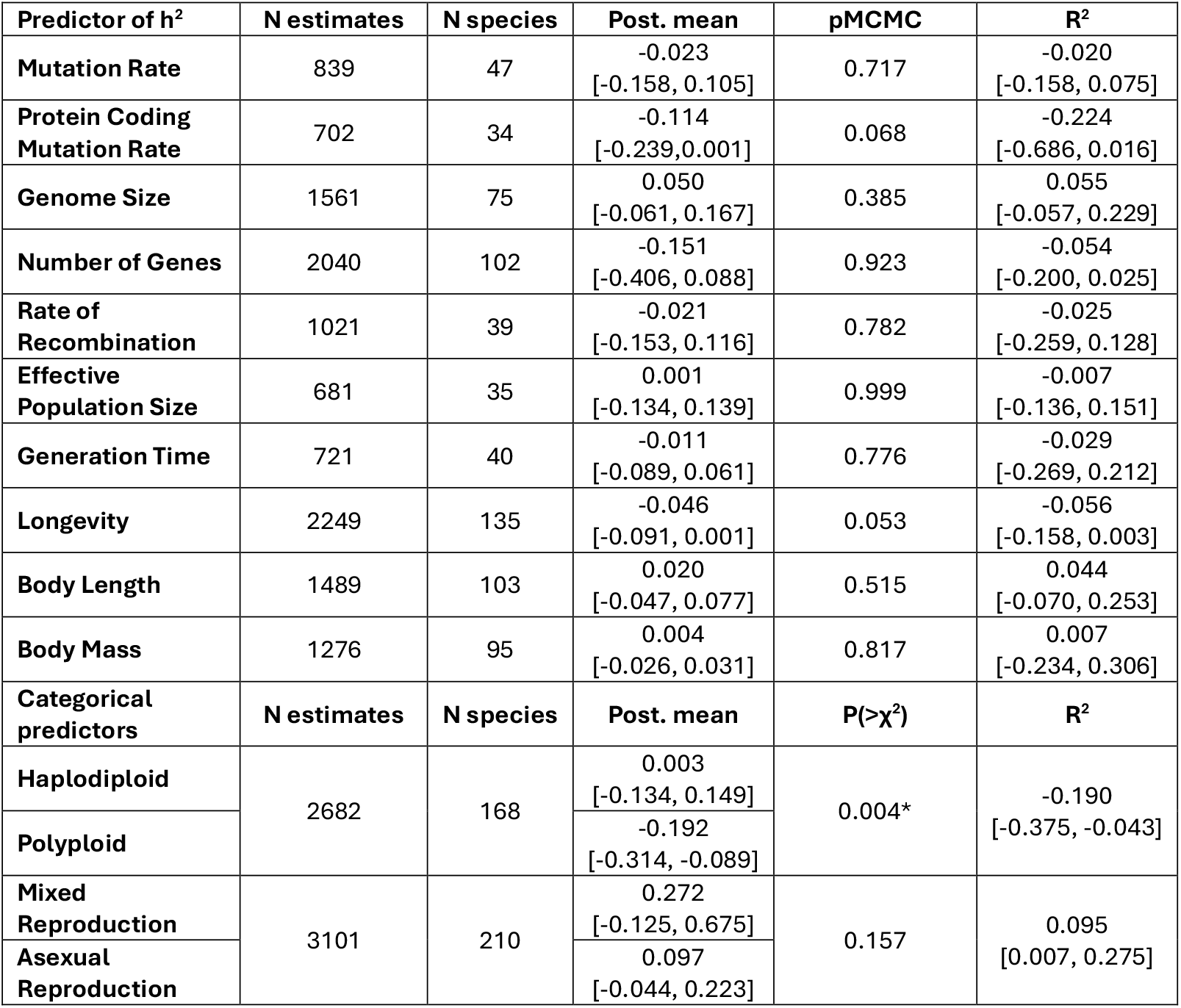
The regression of heritability on various variables. Given is the number of estimates and the number of species from which those estimates come from for each analysis, along with the posterior means with their CIs and p_MCMC_ (twice the posterior probability that the effect lies on the opposite site of zero from its estimated direction) as well as the signed R^2^ for all h^2^ indicating the proportion of the between-species variance explained. For the categorical predictors the p-value of the omnibus test are denoted as P(>X^2^). For the categorical predictors, the posterior means are the deviation from the reference levels (Diploid and Sexual, respectively). An * in the p_MCMC_ or P(>χ^2^) column indicates a significant correlation after Bonferroni correction.

| Predictor of $h^2$ | N estimates | N species | Post. mean | $p_{MCMC}$ | $R^2$ |
| --- | --- | --- | --- | --- | --- |
| **Mutation Rate** | 839 | 47 | -0.023<br>[-0.158, 0.105] | 0.717 | -0.020<br>[-0.158, 0.075] |
| **Protein Coding Mutation Rate** | 702 | 34 | -0.114<br>[-0.239, 0.001] | 0.068 | -0.224<br>[-0.686, 0.016] |
| **Genome Size** | 1561 | 75 | 0.050<br>[-0.061, 0.167] | 0.385 | 0.055<br>[-0.057, 0.229] |
| **Number of Genes** | 2040 | 102 | -0.151<br>[-0.406, 0.088] | 0.923 | -0.054<br>[-0.200, 0.025] |
| **Rate of Recombination** | 1021 | 39 | -0.021<br>[-0.153, 0.116] | 0.782 | -0.025<br>[-0.259, 0.128] |
| **Effective Population Size** | 681 | 35 | 0.001<br>[-0.134, 0.139] | 0.999 | -0.007<br>[-0.136, 0.151] |
| **Generation Time** | 721 | 40 | -0.011<br>[-0.089, 0.061] | 0.776 | -0.029<br>[-0.269, 0.212] |
| **Longevity** | 2249 | 135 | -0.046<br>[-0.091, 0.001] | 0.053 | -0.056<br>[-0.158, 0.003] |
| **Body Length** | 1489 | 103 | 0.020<br>[-0.047, 0.077] | 0.515 | 0.044<br>[-0.070, 0.253] |
| **Body Mass** | 1276 | 95 | 0.004<br>[-0.026, 0.031] | 0.817 | 0.007<br>[-0.234, 0.306] |
| **Categorical predictors** | **N estimates** | **N species** | **Post. mean** | **$P(>\chi^2)$** | **$R^2$** |
| **Haplodiploid** | 2682 | 168 | 0.003<br>[-0.134, 0.149] | 0.004* | -0.190<br>[-0.375, -0.043] |
| **Polyploid** |  |  | -0.192<br>[-0.314, -0.089] |  |  |
| **Mixed Reproduction** | 3101 | 210 | 0.272<br>[-0.125, 0.675] | 0.157 | 0.095<br>[0.007, 0.275] |
| **Asexual Reproduction** |  |  | 0.097<br>[-0.044, 0.223] |  |  |

### Evolvability versus Heritability

Although estimates of I_A_ and h^2^ are significantly correlated across individual estimates, the correlation is weak (r=0.24 [0.20, 0.29]; Figure S7), and there is no significant correlation between I_A_ and h^2^ across species (r=0.13 [-0.13, 0.38]; Figure S7). There are several reasons why I_A_ and h^2^ might be weakly correlated. In particular, it has previously been shown that the residual variance -the sum of the non-additive genetic and environmental variances - is positively correlated to V_A_ across individual estimates (Hansen *et al*., 2011). Consistent with this, we find a strong positive correlation between residual variance (mean-scaled, I_R_) and I_A_ across individual estimates (r=0.73 [0.70, 0.75]; Figure S8) and species (r=0.69 [0.47, 0.89]; Figure S8). Similarly, we find a significantly negative relationship between h^2^ and I_R_ across individual estimates (r=-0.33 [-0.36, - 0.29]; Figure S9) and species (r=-0.51 [-0.68, -0.28]; Figure S9).

In addition to the correlations we also analysed the interspecific variation in I_R_ itself using an interspecific variation model and find that the variation in I_R_ follows a similar pattern to I_A_; there is a strong phylogenetic component (interspecific variation= 29% [8.3%, 53%]) (Table S6) with plants having substantially more residual variance than other phylogenetic groups (Figure S10).

## Discussion

We find significant interspecific variation for two measures of additive genetic variance, the evolvability (I_A_), the additive genetic variance divided by the square of the mean, and the heritability (h^2^). For evolvability a substantial portion of the observed variation can be attributed to phylogenetic relatedness—closely related species tend to exhibit more similar evolvabilities than distantly related species. Much of this phylogenetic signal appears to be driven by plants, which exhibit markedly higher evolvability than animals. Despite this broad taxonomic trend, we also observe substantial variation within major clades, including within both plants and animals, and within most taxonomic groups for which we have sufficient data—birds being the notable exception (Table 2). The variation in evolvability is substantial; we estimate that the top 25% of species have more than 9-fold [3.5, 23] more evolvability than the bottom 25% of species.

An important question is whether the variation we observe in our analysis is due to variation between studies and populations, rather than between species, given that measures of V_A_ can vary between populations of the same species (e.g. Quéméré *et al*., 2018; Muff *et al*., 2019; Windig *et al*., 2004). Several lines of evidence suggest that there is genuine variation between species. In particular, there is a substantial phylogenetic effect for I_A_ that cannot be explained by population-level variation. Moreover, our overall model structure - including study-level and other random effects - effectively accounts for variation among populations within species.

Previously, Young and Postma (2023) found significant variation in h^2^ across mammalian and avian species, without a detectable phylogenetic signal, but no evidence of variation in CV_A_, the square root of I_A_. We performed a comparable analysis using our database of estimates from mammals and birds. Their original dataset had 1822 heritabilities for 68 species and only 378 CV_A_ estimates for 23 species, we compiled 1118 heritabilities for 82 species and 698 estimates of I_A_ for 68 species. We find interspecific variation in evolvability but not in heritability for this dataset. Biologically, our results are broadly concordant with those of Young and Postma (2023), and the apparent differences in findings likely reflect methodological rather than biological factors.

### Why does evolvability vary between species?

Although we find substantial variation in evolvability we do not find that it is predicted by any variable that we have tested except body length. Particularly surprising is the lack of a correlation between evolvability and the mutation rate since most models of the maintenance of the additive genetic variance predict that it should depend on the mutational variance (Walsh and Lynch, 2018). There are several reasons why we might not have detected an effect. First, available proxies for mutational input are relatively crude. We assessed whether evolvability correlates with the nucleotide mutation rate alone, as well as with the mutation rate scaled by proteome size; for these variables we have few estimates. We also considered the number of protein coding genes (without multiplying this by the mutation rate, which is only known for a limited number of species). However, the number of genes likely represents a limited proxy for the genomic regions in which mutations can generate phenotypic effects. It does not account for the amount of regulatory DNA, which is likely to vary among species. Moreover, under most models of quantitative genetic variation maintenance mutational variance is influenced not only by mutation rate but also by the distribution of effect sizes, which is also known to vary across species (Huber *et al*., 2017; Castellano *et al*., 2019). Unfortunately, we have very few estimates for the distribution of effect sizes. However, possibly the biggest omission in our estimate of the mutation rate are structural variants (SVs) including transposable element (TE) insertions. There is evidence from Drosophila (Long *et al*., 2000; Robillard *et al*., 2016) and yeast (Loegler *et al*., 2025) that SVs can contribute substantially to additive genetic variation. Unfortunately, estimates for the rate at which SVs occur have been estimated for very few species. It is also possible that I_A_ is not correlated to our estimates of the mutational variance because it does not differ substantially between species, in which case it could not account for the variation in I_A_ we observe.

We also find no correlation between evolvability and the effective population size. This might be due to a lack of power; both our estimates of N_e_ and species mean I_A_ are subject to reasonable levels of error. However, it might reflect the fact that most of the mutational input to the additive genetic variance is deleterious and subject to strong selection; under such a model the genetic variance is likely to be independent of N_e_ unless N_e_ is very small (Keightley and Hill, 1988; Keightley and Hill, 1990; Bürger *et al*., 1989). It might also be that additive genetic variance is maintained by balancing selection (Walsh and Lynch, 2018).

The one significant correlation we detect is a positive relationship between evolvability and body size in animals (measured as the maximum linear dimension), which remains statistically significant after correcting for multiple tests (p = 0.003, multiple-test–corrected significance threshold=0.004). This might arise because larger bodied species tend to have longer generation times, and species with long generation times have higher mutation rates per generation (Bergeron *et al*., 2023; Wang and Obbard, 2023) although we tested for a correlation between evolvability and the mutation rate we had few estimates in our analysis and hence low power. There is also some evidence that larger bodied species have larger ranges (Pyron, 2002; Newsome *et al*., 2019) and might therefore be exposed to different environments, leading to maintenance of locally adaptive genetic variation. Finally, the observed positive correlation may also be influenced by trait scaling. If the additive genetic variance does not increase linearly with the square of the trait mean, I_A_ will tend to increase with larger trait values. While this could in principle be tested by excluding size traits, the majority of our dataset consists of such traits, and removing them would result in a severe loss of power, preventing reliable inference.

Although this study explored a range of factors that could explain interspecific variation in quantitative genetic parameters, there are additional important forces we have not explored. One such factor is selection, which is expected to affect levels of additive genetic variance (though not always under directional selection - see Hill, 1982). However, the extent to which selection varies among species is largely unknown, because meta-analyses of selection gradients rarely account for species identity or phylogenetic relatedness (Hereford *et al*., 2004). Finally, population structure - such as sub-division and migration - could influence V_A_, but we have little information on these for the species we have analysed.

### Why is evolvability higher in plants?

We find that plants have significantly higher evolvability than animals. Although, our analysis suggests that the difference between plants and animals is no more than we might expect under a simple phylogenetic model of trait evolution, the difference is substantial, with plants estimated to have ∼4.7-fold [2.7, 8.3] more evolvability than animals.

Unfortunately, understanding why plants and animals differ is challenging because there are many fundamental differences between plants and animals, and both groups are monophyletic; as a result, we have limited statistical power to test hypotheses. For example, plants generally have many more genes than animals, but because there is relatively little variation in gene number within plants and animals, we have few independent contrasts to detect an association. We find no evidence that this pattern might be attributable to differences in the mutation rate, effective population size, mating system, genome size or ploidy. However, there are two conspicuous differences between plants and animals that might contribute to the difference in evolvability between the groups. First, plants have an additional genome, in the chloroplast. While both plastid and mitochondrial genomes are small relative to the nuclear genome, studies have shown that genetic variation in the mitochondrial genome can significantly contribute to phenotypic variation in animals (Dowling *et al*., 2007; Clancy, 2008). A meta-analysis of studies in which mtDNA was introgressed between nuclear backgrounds in animals suggested the effects can be substantial, with nearly 50% of cases changing the mean up or down by >10% (Eyre-Walker, 2017) although this is likely to be an overestimate due to sampling error (Morrissey, 2016). Surprisingly, little is known about the genetic diversity contained within the plastid genome and whether it contributes to QGV. There are some species for which plastid-nuclear incompatibilities have been shown to be segregating (Postel *et al*., 2022), but little work has been done beyond this.

The second major difference between plants and animals is the potential for transgenerational epigenetic inheritance (TEI) (Bond and Baulcombe, 2014). There are examples of TEI in both plants and animals (Bošković and Rando, 2018) but no comparative study of how much epigenetic variation is inherited in the two groups of organisms. However, there are some reasons for believing that inherited epigenetic variation might be greater in plants than animals. There are several pathways by which TEI can be propagated (Bošković and Rando, 2018) but the simplest is through DNA methylation, and most methylation marks are not removed during gametogenesis in plants, whereas extensive remodelling occurs in animals (Bošković and Rando, 2018; Bond and Baulcombe, 2014). Experiments using epigenetic recombinant inbred lines (epiRILs) of *Arabidopsis thaliana* suggest that heritable differences in methylation can generate substantial phenotypic variation (Reinders *et al*., 2009; Roux *et al*., 2011) comparable in magnitude to differences observed among natural accessions. Broad sense heritability amongst epiRILs can be as high as 90% for some traits, such as flowering time, but essentially absent for others (Roux et al., 2011; Postel *et al*., 2022). However, an epiRIL experiment gives an upper estimate for the variation induced by heritable methylation since it involves crossing methylated and largely unmethylated parental strains, hence generating more diversity in methylation than would be seen in a natural population. The fact that largely unmethylated strains are viable suggests that DNA methylation might not play an essential or widespread role in generating large amounts of additive genetic variance, even if it can influence particular phenotypes. Although natural populations do contain epigenetic variation in methylation (Schmitz *et al*., 2013; Dubin *et al*., 2015), much of this appears to be genetically determined (Dubin *et al*., 2015). It therefore seems unlikely that epigenetic variation is why plants have higher evolvability than animals.

### Evolvability - other effects

Hansen *et al*. (2011) reported that the evolvability of cubic traits was roughly nine times that of linear traits, implying strong genetic correlations among trait dimensions (e.g., between height and width). In contrast, our analysis finds that the ratio of linear, quadratic, and cubic traits is closer to 1:2:3, consistent with variation in different dimensions being largely independent. The difference between our study and that of Hansen *et al*. (2011) might be due to our larger sample size and the fact that we have modelled many effects they did not, including the method of estimation, trait type and the relatedness of the species in our sample; if we simply take the median estimates of the observed I_A_ values we find a pattern closer to that originally reported by Hansen *et al*. (2011). An additional difference is that our analyses are based on log-transformed I_A_ values, whereas Hansen *et al*., (2011) used raw I_A_, which will also contribute to the differences we observe. Biologically, one might have expected an intermediate scaling between the different dimensions, given that our dataset is extremely heterogeneous spanning diverse species and traits that likely vary in their degree of dimensional integration. However, a more robust test of these scaling relationships will require a dataset with a greater representation of higher-dimensional traits.

We also show, for the first time, that count traits exhibit significantly higher evolvability (I_A_) and lower heritability (h^2^) compared to linear traits. While Hansen *et al*. (2011) did consider counts they did so only for size traits. The relatively high I_A_ of count traits may reflect a combination of biological and statistical factors. Biologically, count traits often arise from modular or discrete developmental processes, where variation among semi-independent units (e.g. number of leaves or offspring) can accumulate additively. Methodologically, the scaling of variance by the mean may contribute to elevated I_A_ values when counts are low or distributions are skewed, as we might expect under a Poisson distribution, overdispersion or zero-inflated distributions, although not all count traits conform to such distributions.

Interestingly, when we examine evolvability across different trait types we find no significant difference between life-history and morphological traits, contrary to previous analyses (Hansen *et al*., 2011; Young and Postma, 2023). Our analysis however varies from previous studies in several ways. Firstly, we separate fitness and life-history, yet neither is significantly different from morphological traits in I_A_. We also include a random effect grouping traits that, although labelled differently across publications, measure the same underlying property (e.g. body size). Lastly, we explicitly account for trait dimensions across all trait types. If we remove trait dimension from our analysis, we find that life-history traits have higher evolvability than morphology traits. This result is particularly interesting when considering the distribution of trait dimensions across the different trait types. Morphological traits are predominantly linear (65%), with smaller proportions being quadratic (5.3%) or cubic (21%). In contrast life-history traits are mostly durations (46%) and counts (51%). These distributions suggest that apparent differences between life-history and morphological traits may reflect underlying differences in trait dimensionality rather than intrinsic differences in evolvability.

Hansen *et al*. (2011) demonstrated that the residual variance (the sum of the environmental and non-additive genetic variance) is positively correlated with the additive genetic variance across individual estimates of these quantities, a pattern we replicate. They argue that the relationship exists because increases in allele frequencies are expected to increase the additive and epistatic variances (Hansen and Wagner 2001). Furthermore, the additive, epistatic and dominance variances all depend on the number of loci underlying a trait. Houle (1992) has also argued that the environmental variance should increase with the additive genetic variance across trait types because complex traits are probably more sensitive to both genetic and environmental perturbation.

Extending these findings, we find a strong and significant relationship between I_R_ and I_A_ across species, demonstrating that the positive relationship observed at the level of individual estimates also holds at the species level. This is consistent with our observation that interspecific variation in I_R_ follows the same trend as I_A_ – phylogenetic signal with higher estimates in plants. This suggests that factors linking the additive and residual variances operate not only within species but also across species.

### Heritability

We find significant variation in heritability across species, with the upper and lower quartiles varying by nearly two-fold. This variation could reflect differences in additive, non-additive or environmental variance. Across species, we find no evidence of a relationship between I_A_ and h^2^, suggesting that the interspecific variation in h^2^ are unlikely to be driven primarily by differences in the additive genetic variance. In contrast, the mean-scaled residual variance (I_R_) varies significantly between species and is strongly positively correlated with I_A_ and negatively correlated with h^2^ across both estimates and species. This pattern suggests, as proposed by Young and Postma (2023), that the observed between species variation in h^2^ is likely influenced more by differences in residual variance than by differences in the additive genetic variance.

As with evolvability, we find only one variable that explains some of the variation in h^2^, this is ploidy level. Polyploids exhibit significantly lower h^2^ than diploids, which may reflect increased opportunities for epistasis and other non-additive genetic interactions in polyploids. Some of this epistatic variance may contribute to the additive genetic variance depending on the experimental design, but overall, it is consistent with the idea that higher levels of non-additive variation obscure the relationship between I_A_ and h^2^ at the species level We find evidence that estimates of h^2^ are generally higher in the lab than in natural populations contrary to a previous meta-analysis (Weisenberg and Roff, 1996). However, we have many more estimates and probably increased power because we have modelled many fixed effects that influence h^2^. The difference between lab and wild estimates is not likely to be due to higher V_A_ in lab populations, as I_A_ is not significantly elevated in lab populations. A commonly invoked explanation is that laboratory conditions reduce environmental variance, thereby inflating heritability estimates. However, we find no significant difference between lab and natural populations in I_R_, the residual variance, which is the sum of the non-additive genetic and environmental variances. This suggests that the higher heritabilities observed in laboratory studies cannot be attributed simply to reduced residual variance under controlled environments and may reflect more subtle differences between laboratory and natural settings that affect how genetic and environmental variation contribute to phenotypic variance.

Consistent with previous literature, we find that morphological traits have higher heritability than other trait types (e.g. Young and Postma, 2023; Hansen *et al*., 2011). In contrast, count traits exhibit lower h^2^ despite having high I_A_. This pattern mirrors our findings that count traits have significantly higher I_R_ (Table S6). Because heritability is sensitive to the residual variance whereas I_A_ is not, elevated I_R_ in count traits can lead to reduced h^2^ even when evolvability is high.

### Summary

We find substantial interspecific variation in evolvability (I_A_), with clear phylogenetic signal indicating that closely related species tend to have more similar levels of evolvability, with plants as sampled here standing out with particularly7 high levels. While we also find significant interspecific variation in heritability (h^2^) the absence of a correlation with I_A_ suggests that this variation might be due to variation in the residual variance. Although we find significant variation in the additive genetic variance between species, we are unable to explain this. Nevertheless, our results suggest that some species have a substantially greater capacity to evolve.

## Acknowledgements

We are very grateful to Maria Clara Castellanos, Pierre Nouvellet, Bill Hughes and the rest of the University of Sussex evolutionary genetics community for helpful discussion. We also thank Thomas Hansen, David Houle and Christophe Pelabon for the valuable and helpful discussion on estimates of evolvability. L.Z. is funded by the John Maynard Smith PhD Studentship.

