## Supplementary Material for "Levels of additive genetic variation vary substantially between species"

**Table S2: Data Summary**

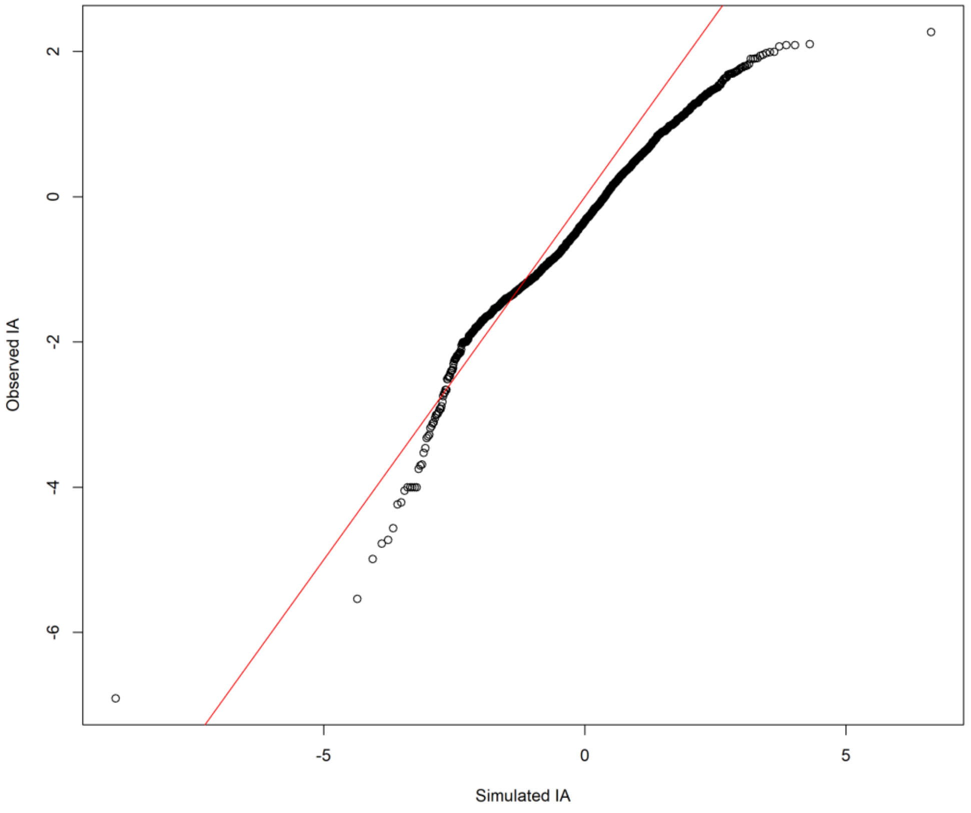


**Figure S1:** QQplot for visual inspection of model fit. Simulated $I_A$ values using the simulate() function plotted in a qqplot against the observed $I_A$ values.

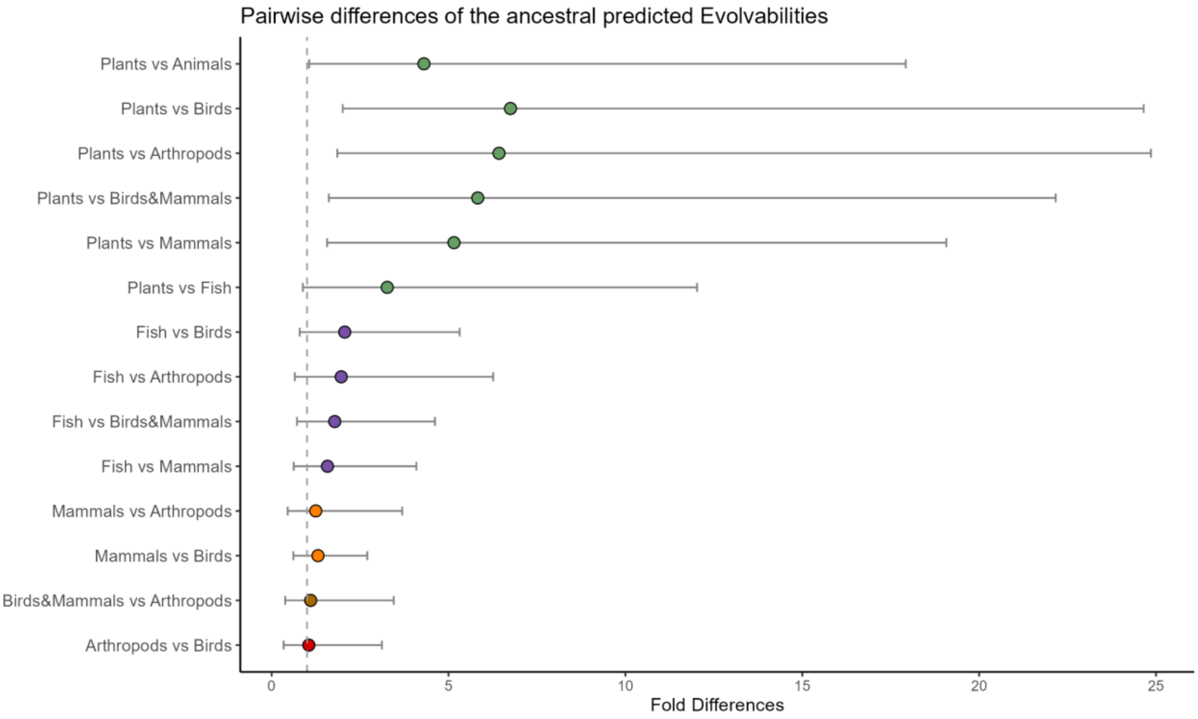


**Figure S2:** The difference in ancestral evolvability between groups of animals and plants. Each point represents the posterior mean difference between two ancestral nodes of the phylogeny, back-transformed to the original data scale. Horizontal bars indicate the 95% credible intervals. The dashed line at 1 indicates no difference between groups. Differences greater than 1 indicate that the first node (first-mentioned group in the y-axis) has a x-fold higher predicted evolvability than the second node. All comparisons are plotted so that the means are positive.

**Table S3:** Summary of the random and fixed effects for the interspecific variation model of evolvability with the inclusion of the taxonomic (Plant/Animal) fixed effect. Random effects as the proportion of variance explained by them and the p-values of an omnibus test for the fixed effects.

| **Random Effect** | **mean** | **l-95%** | **u-95%** | **Percentage variance explained** |
|---|---|---|---|---|
| **Phylogeny** | 0.590 | 0.105 | 1.143 | 40% [17, 63] |
| **Non-Phylogenetic** | 0.012 | 2e-09 | 0.048 | 1.0% [0, 4.0] |
| **Publication_ID** | 0.182 | 0.118 | 0.253 | 14% [7.5, 22] |
| **Trait_2** | 0.176 | 0.121 | 0.233 | 13% [7.1, 19] |
| **Residual** | 0.424 | 0.391 | 0.462 | 32% [21, 45] |

| | **mean** | **l-95%** | **u-95%** | **pMCMC** | **P(>$x^2$)** |
|---|---|---|---|---|---|
| **Intercept** | -0.769 | -1.899 | 0.534 | 0.216 | |
| **Method: clonal** | -0.092 | -0.534 | 0.330 | 0.688 | 0.011 |
| **Method: full-sib** | 0.611 | 0.226 | 1.030 | 0.003 | |
| **Method: half-sib** | 0.033 | -0.203 | 0.286 | 0.810 | |
| **Method: mid-parent-offspring** | -0.401 | -0.782 | -0.034 | 0.040 | |
| **Method: realized** | -0.478 | -2.653 | 1.417 | 0.643 | |

| | | | | | |
|---|---|---|---|---|---|
| **Method: single-parent-offspring** | 0.088 | -0.192 | 0.367 | 0.553 | |
| **Trait type: behaviour** | 0.728 | 0.404 | 1.048 | <4e-04 | <4e-04 |
| **Trait type: fitness** | -0.071 | -0.700 | 0.604 | 0.842 | |
| **Trait type: life history** | 0.027 | -0.238 | 0.305 | 0.838 | |
| **Trait type: physiology** | 0.190 | -0.031 | 0.453 | 0.123 | |
| **Dimension: quadratic** | 0.220 | -0.076 | 0.532 | 0.155 | <4e-04 |
| **Dimension: cubic** | 0.544 | 0.356 | 0.764 | <4e-04 | |
| **Dimension: time** | 0.233 | -0.072 | 0.552 | 0.148 | |
| **Dimension: count** | 0.531 | 0.297 | 0.779 | <4e-04 | |
| **Dimension: other** | 0.270 | 0.024 | 0.520 | 0.031 | |
| **n_fixed** | 0.010 | -0.029 | 0.050 | 0.642 | 0.645 |
| **n_random** | 0.093 | 0.004 | 0.179 | 0.036 | 0.038 |
| **Environment: lab** | 0.167 | -0.086 | 0.427 | 0.213 | 0.002 |
| **Environment: field/lab** | -0.324 | -0.671 | 0.002 | 0.062 | |
| **Taxonomic Group: Plants** | 0.729 | -1.456 | 2.520 | 0.417 | 0.453 |

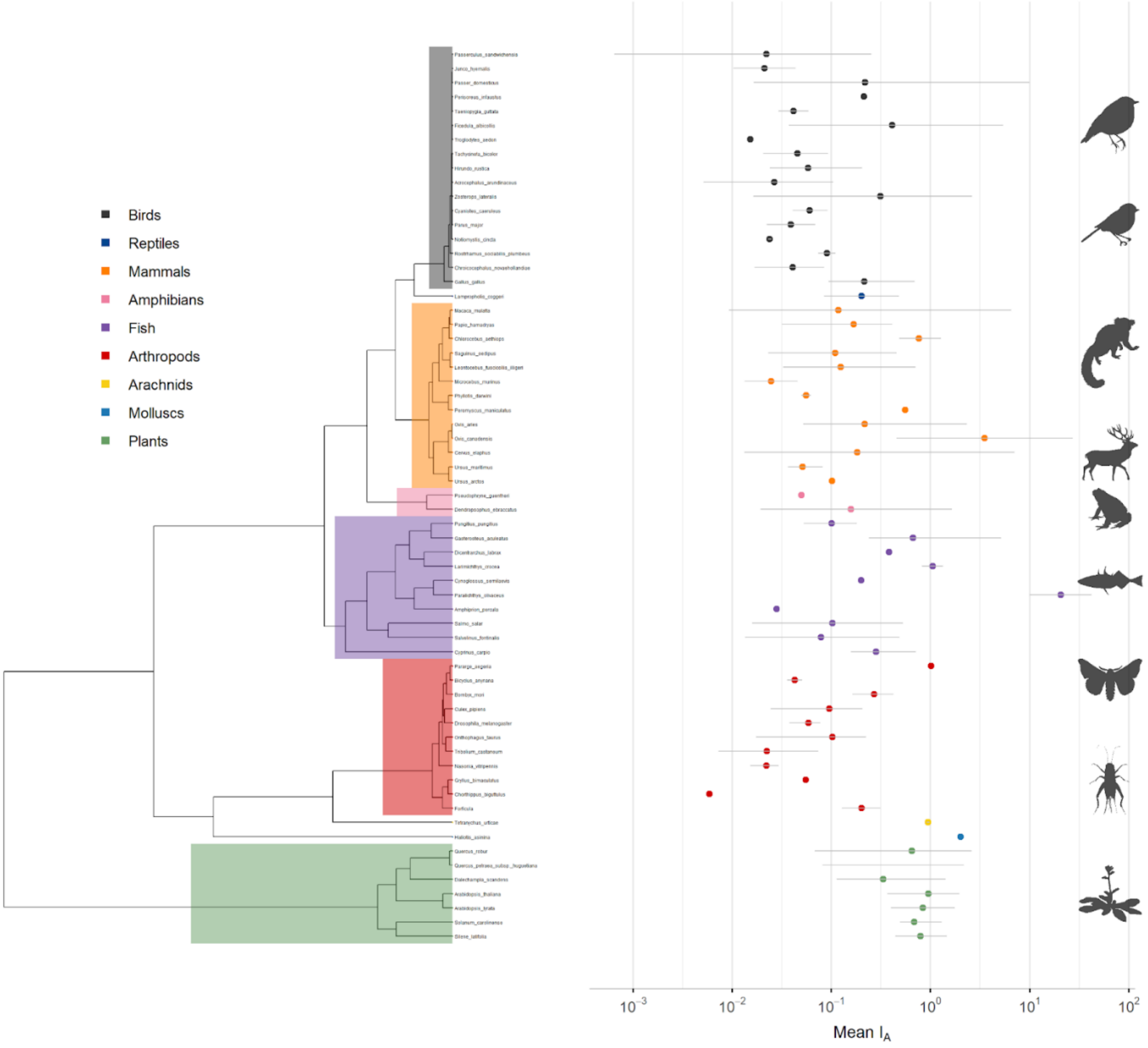


**Figure S3:** Mean $I_A$ for each species for linear morphological traits which were estimated using an animal model.

**Table S4:** Summary of the random and fixed effects for the interspecific variation model of evolvability. Random effects as the proportion of variance explained by them and the p-values of an omnibus test for the fixed effects.

| **Random Effect** | **mean** | **l-95%** | **u-95%** | **Percentage variance explained** |
|---|---|---|---|---|
| **Phylogeny** | 0.530 | 0.135 | 1.011 | 38% [15, 58] |
| **Non-Phylognetic** | 0.012 | 8e-09 | 0.045 | 1.0% [0, 3.9] |
| **Publication_ID** | 0.182 | 0.123 | 0.251 | 14% [7.8, 22] |
| **Trait_2** | 0.177 | 0.123 | 0.233 | 14% [7.9, 20] |
| **Residual** | 0.423 | 0.388 | 0.459 | 33% [23, 45] |

| | **mean** | **l-95%** | **u-95%** | **pMCMC** | **$P(>x^2)$** |
|---|---|---|---|---|---|
| **Intercept** | -0.443 | -1.294 | 0.470 | 0.274 | |
| **Method: clonal** | -0.100 | -0.536 | 0.367 | 0.643 | 0.012 |
| **Method: full-sib** | 0.601 | 0.195 | 0.987 | 0.005 | |
| **Method: half-sib** | 0.037 | -0.220 | 0.276 | 0.790 | |
| **Method: mid-parent-offspring** | -0.404 | -0.771 | -0.034 | 0.035 | |
| **Method: realized** | -0.503 | -2.757 | 1.490 | 0.623 | |
| **Method: single-parent-offspring** | 0.092 | -0.185 | 0.383 | 0.528 | |
| **Trait type: behaviour** | 0.717 | 0.388 | 1.025 | <4e-04 | <4e-04 |
| **Trait type: fitness** | -0.055 | -0.748 | 0.555 | 0.866 | |
| **Trait type: life history** | 0.028 | -0.238 | 0.313 | 0.840 | |
| **Trait type: physiology** | 0.191 | -0.071 | 0.415 | 0.140 | |
| **Dimension: quadratic** | 0.223 | -0.082 | 0.544 | 0.173 | <4e-04 |
| **Dimension: cubic** | 0.541 | 0.335 | 0.740 | <4e-04 | |
| **Dimension: time** | 0.234 | -0.108 | 0.542 | 0.143 | |
| **Dimension: count** | 0.531 | 0.277 | 0.772 | <4e-04 | |
| **Dimension: other** | 0.267 | 0.009 | 0.519 | 0.043 | |
| **n_fixed** | 0.010 | -0.029 | 0.048 | 0.600 | 0.601 |
| **n_random** | 0.092 | 0.007 | 0.177 | 0.033 | 0.040 |
| **Environment: lab** | 0.172 | -0.078 | 0.414 | 0.162 | 0.003 |
| **Environment: field/lab** | -0.305 | -0.644 | 0.035 | 0.080 | |

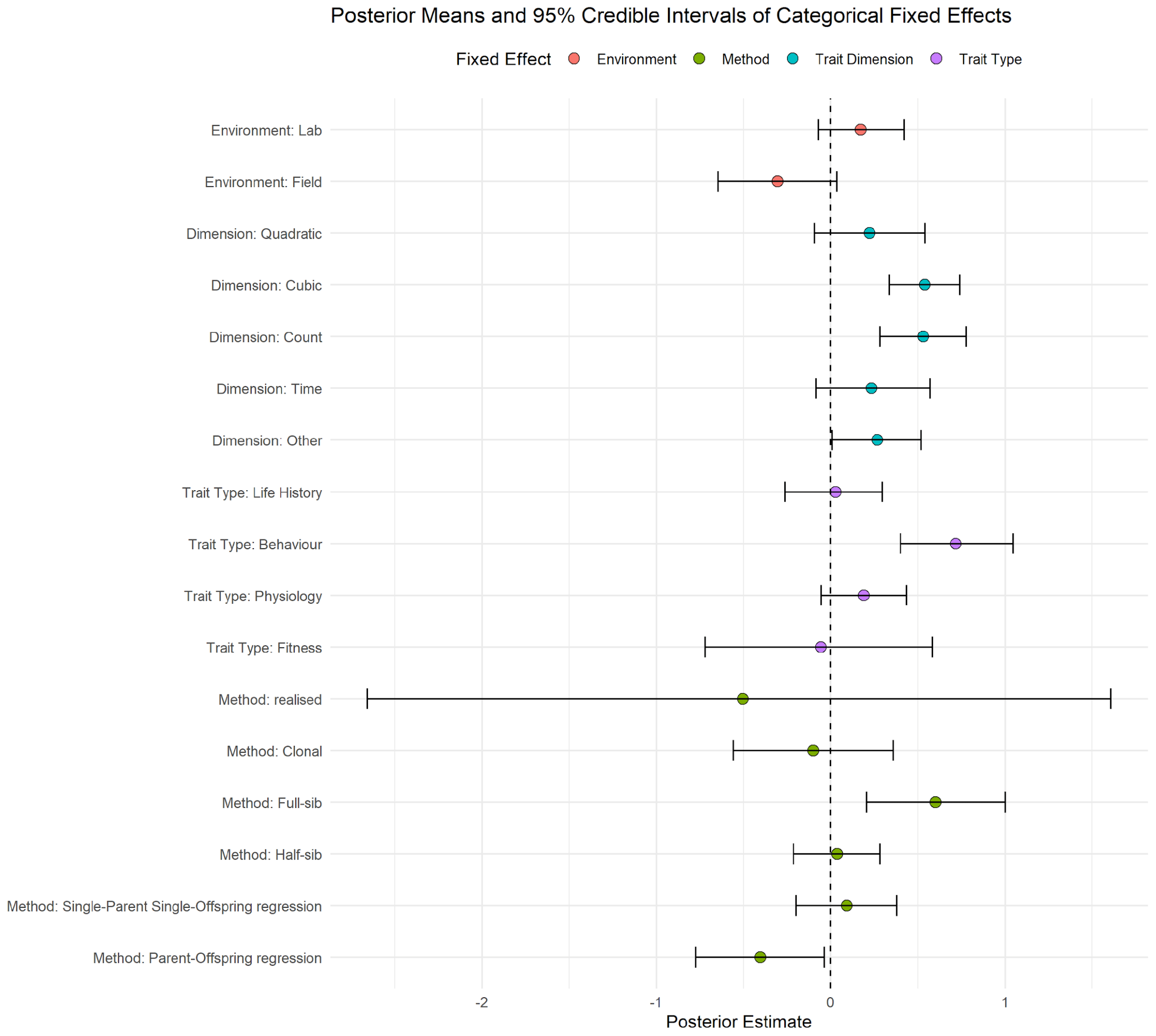


**Figure S4:** The posterior estimate for each fixed effect in the statistical model of $I_A$, along with its 95% credibility intervals; all estimates are deviations from the default model of a linear morphological trait estimated using an animal model from a population in the wild.

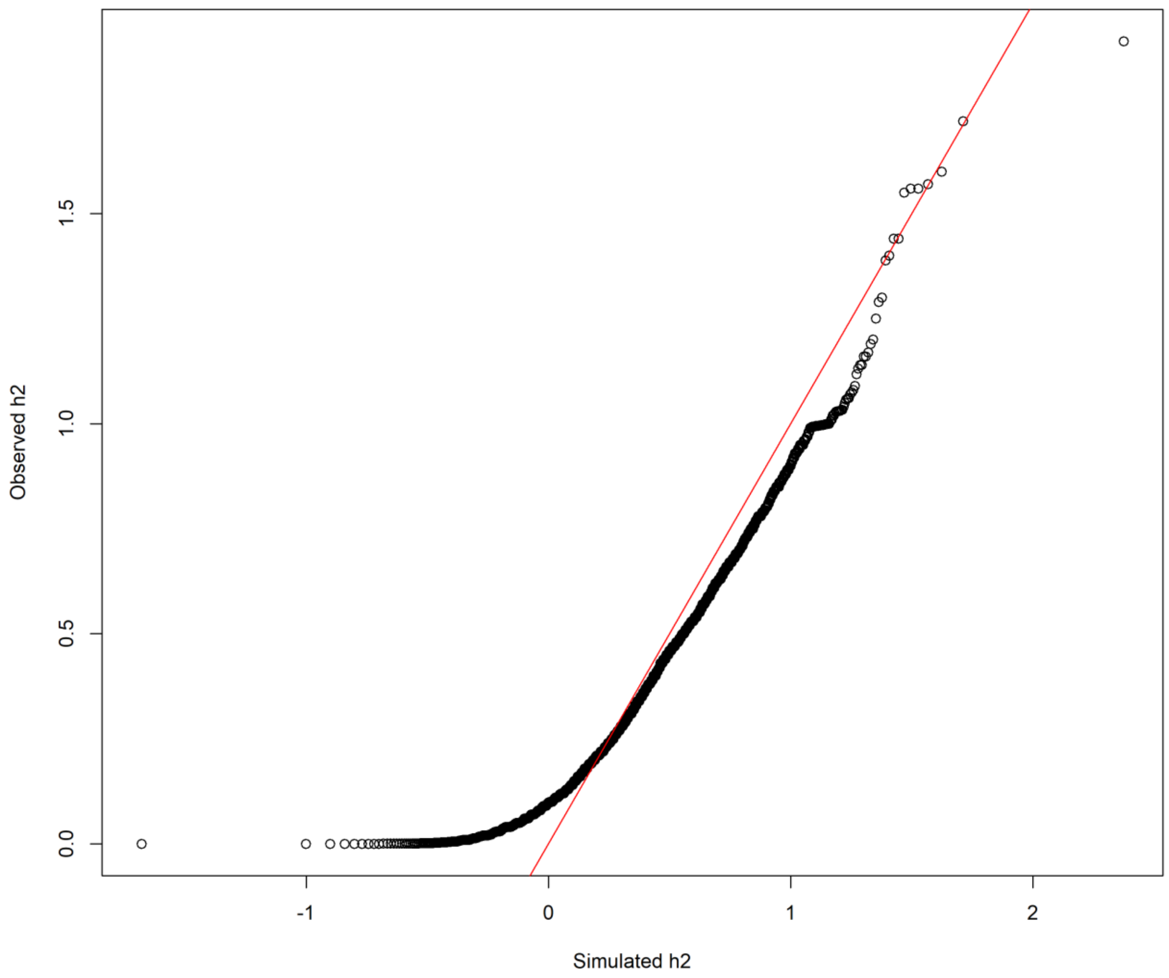


**Figure S5:** QQplot for visual inspection of model fit. Simulated $h^2$ values using the simulate() function plotted in a qqplot against the observed $h^2$ values.

**Table S5:** Summary of the random and fixed effects for the interspecific variation model of heritability. random effects as the proportion of variance explained by them and the p-values of an omnibus test for the fixed effects.

| Random Effect | mean | l-95% | u-95% | Percentage variance explained |
|---|---|---|---|---|
| **Phylogeny** | 0.034 | 4e-09 | 0.090 | 28% [0.35, 57] |
| **Non-Phylogenetic** | 0.004 | 1e-08 | 0.010 | 4.4% [0, 12] |
| **Publication_ID** | 0.023 | 0.017 | 0.029 | 22% [11, 31] |
| **Trait_2** | 0.004 | 0.002 | 0.006 | 4.2% [1.7, 6.8] |
| **Residual** | 0.042 | 0.039 | 0.044 | 41% [25, 55] |

| | mean | l-95% | u-95% | pMCMC | $P(>x^2)$ |
|---|---|---|---|---|---|
| **Intercept** | 0.326 | 0.110 | 0.557 | 0.013 | |
| **Method: clonal** | -0.026 | -0.136 | 0.092 | 0.668 | <4e-04 |
| **Method: full-sib** | 0.207 | 0.108 | 0.300 | <4e-04 | |
| **Method: half-sib** | -0.004 | -0.075 | 0.063 | 0.906 | |
| **Method: mid-parent-offspring** | 0.070 | -0.013 | 0.149 | 0.094 | |
| **Method: realized** | 0.075 | -0.492 | 0.621 | 0.799 | |
| **Method: single-parent-offspring** | 0.091 | 0.030 | 0.158 | 0.007 | |
| **Trait type: behaviour** | -0.101 | -0.159 | -0.032 | 0.003 | <4e-04 |
| **Trait type: fitness** | -0.147 | -0.284 | -0.013 | 0.033 | |
| **Trait type: life history** | -0.038 | -0.081 | 0.007 | 0.087 | |
| **Trait type: physiology** | -0.075 | -0.117 | -0.032 | 0.001 | |
| **Dimension: quadratic** | -0.015 | -0.088 | 0.054 | 0.684 | 0.012 |
| **Dimension: cubic** | -0.015 | -0.056 | 0.030 | 0.495 | |
| **Dimension: time** | -0.017 | -0.071 | 0.040 | 0.533 | |
| **Dimension: count** | -0.053 | -0.100 | -0.001 | 0.042 | |
| **Dimension: other** | -0.060 | -0.094 | -0.027 | 0.001 | |
| **n_fixed** | -0.008 | -0.018 | 0.002 | 0.118 | 0.115 |
| **n_random** | -0.023 | -0.043 | -0.003 | 0.026 | 0.025 |
| **Environment: lab** | 0.111 | 0.042 | 0.179 | 0.003 | 0.002 |
| **Environment: field/lab** | 0.036 | -0.050 | 0.118 | 0.401 | |

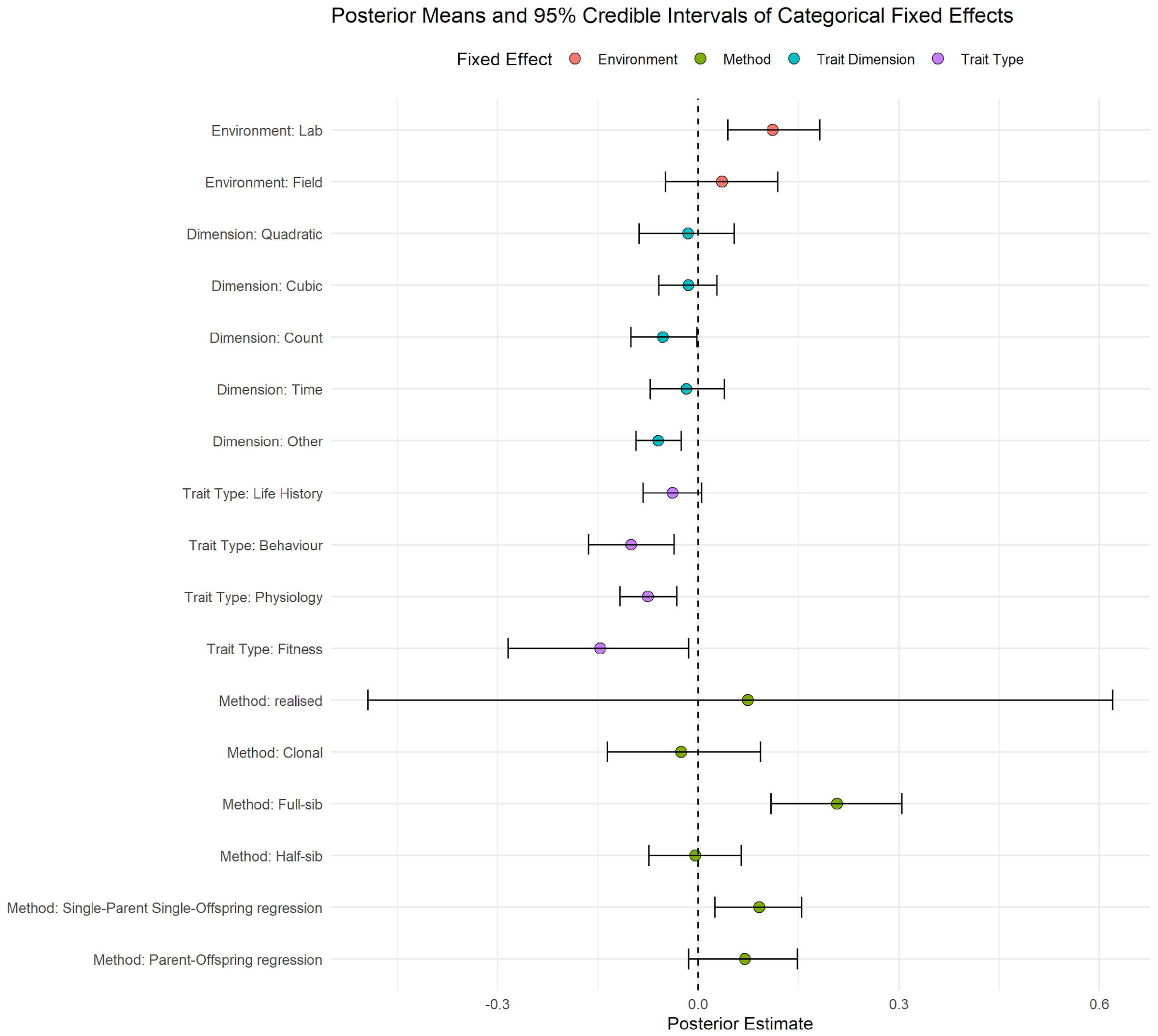


**Figure S6:** The posterior estimate for each fixed effect in the statistical model of $h^2$, along with its 95% credibility intervals; all estimates are deviations from the default model of a linear morphological trait estimated using an animal model from a population in the wild.

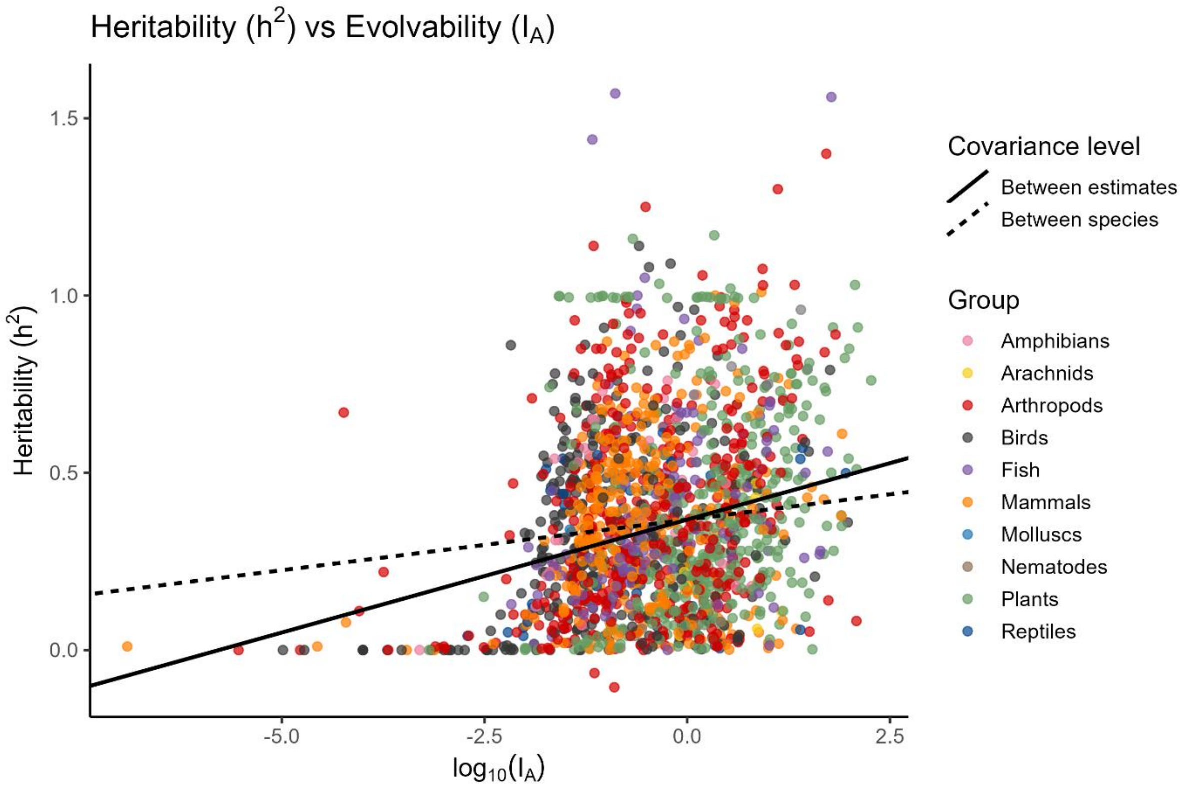


**Figure S7:** The relationship between evolvability ($I_A$) and heritability ($h^2$). Points show observed estimates by taxonomic group. Lines represent slopes derived from different variance-covariance components of the bivariate PGLMM. The between species slope (dashed – phylogenetic +non-phylogenetic) and the between estimate (solid – residual) slope. Lines are anchored at the posterior mean heritability from the bivariate model; intercepts are shown for visual comparison only.

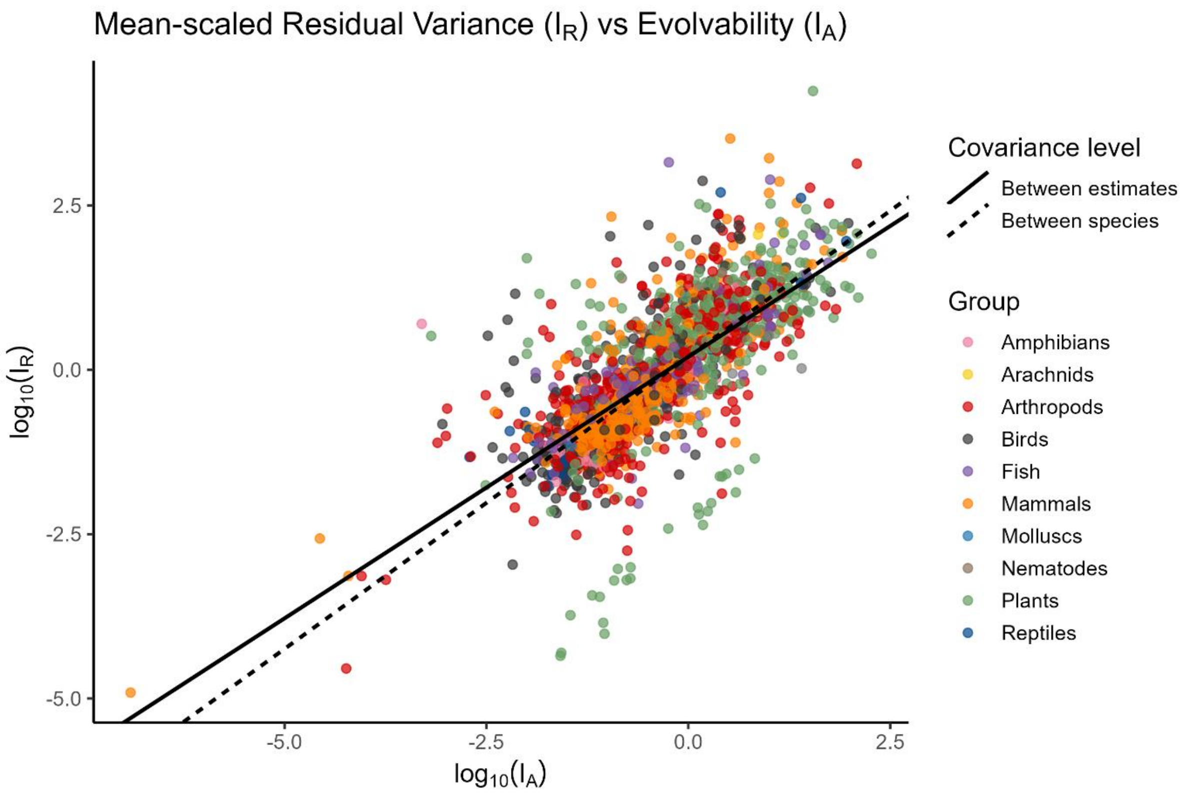


**Figure S8:** The relationship between evolvability ($I_A$) and the mean-scaled residual variance ($I_R$). Points show observed estimates by taxonomic group. Lines represent slopes derived from different variance-covariance components of the bivariate PGLMM. The between species slope (dashed – phylogenetic +non-phylogenetic) and the between estimate (solid – residual) slope. Lines are anchored at the posterior mean heritability from the bivariate model; intercepts are shown for visual comparison only.

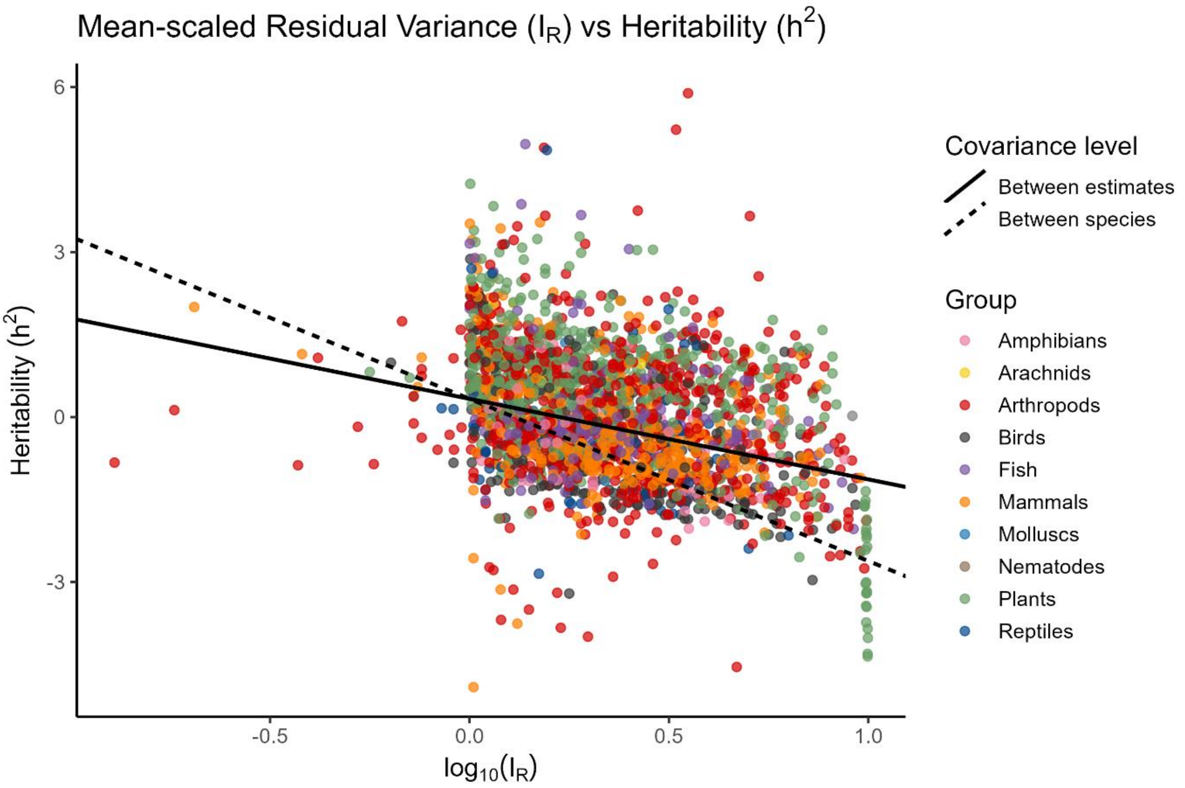


**Figure S9:** The relationship between heritability ($h^2$) and the mean-scaled residual variance ($I_R$). Points show observed estimates by taxonomic group. Lines represent slopes derived from different variance-covariance components of the bivariate PGLMM. The between species slope (dashed – phylogenetic +non-phylogenetic) and the between estimate (solid – residual) slope. Lines are anchored at the posterior mean heritability from the bivariate model; intercepts are shown for visual comparison only.

**Table S6:** Summary of the random and fixed effects for the interspecific variation model of the mean scaled residual variance ($I_R$). Random effects as the proportion of variance explained by them and the p-values of an omnibus test for the fixed effects.

| Random Effect | mean | l-95% | u-95% | Percentage variance explained |
|---|---|---|---|---|
| Phylogeny | 0.291 | 0.018 | 0.662 | 27% [5.9, 51] |
| Non-Phylogenetic | 0.015 | 2e-08 | 0.054 | 1.6% [0, 5.5] |
| Publication_ID | 0.249 | 0.176 | 0.329 | 26% [15, 37] |
| Trait_2 | 0.236 | 0.190 | 0.288 | 24% [16, 32] |
| Residual | 0.213 | 0.193 | 0.233 | 22% [14, 28] |

| | mean | l-95% | u-95% | pMCMC | $P(>x^2)$ |
|---|---|---|---|---|---|
| Intercept | -0.144 | -0.798 | 0.553 | 0.634 | |
| Method: clonal | -0.100 | -0.589 | 0.338 | 0.672 | 0.101 |
| Method: full-sib | 0.303 | -0.166 | 0.766 | 0.200 | |
| Method: half-sib | -0.070 | -0.350 | 0.177 | 0.610 | |
| Method: mid-parent-offspring | -0.470 | -0.859 | -0.113 | 0.017 | |
| Method: realized | -0.848 | -2.594 | 1.040 | 0.359 | |
| Method: single-parent-offspring | -0.240 | -0.550 | 0.039 | 0.104 | |
| Trait type: behaviour | 1.079 | 0.755 | 1.383 | <4e-04 | <4e-04 |
| Trait type: fitness | 1.045 | 0.445 | 1.636 | <4e-04 | |
| Trait type: life history | 0.354 | 0.086 | 0.615 | 0.007 | |
| Trait type: physiology | 0.395 | 0.172 | 0.634 | <4e-04 | |
| Dimension: quadratic | 0.281 | -0.026 | 0.568 | 0.072 | <4e-04 |
| Dimension: cubic | 0.620 | 0.415 | 0.832 | <4e-04 | |
| Dimension: time | 0.350 | 0.032 | 0.636 | 0.033 | |
| Dimension: count | 0.585 | 0.353 | 0.828 | <4e-04 | |
| Dimension: other | 0.352 | 0.104 | 0.601 | 0.008 | |
| n_fixed | -0.005 | -0.047 | 0.039 | 0.800 | 0.812 |
| n_random | 0.125 | 0.025 | 0.219 | 0.010 | 0.014 |
| Environment: lab | -0.166 | -0.491 | 0.180 | 0.331 | 0.624 |
| Environment: field/lab | -0.053 | -0.307 | 0.197 | 0.665 | |

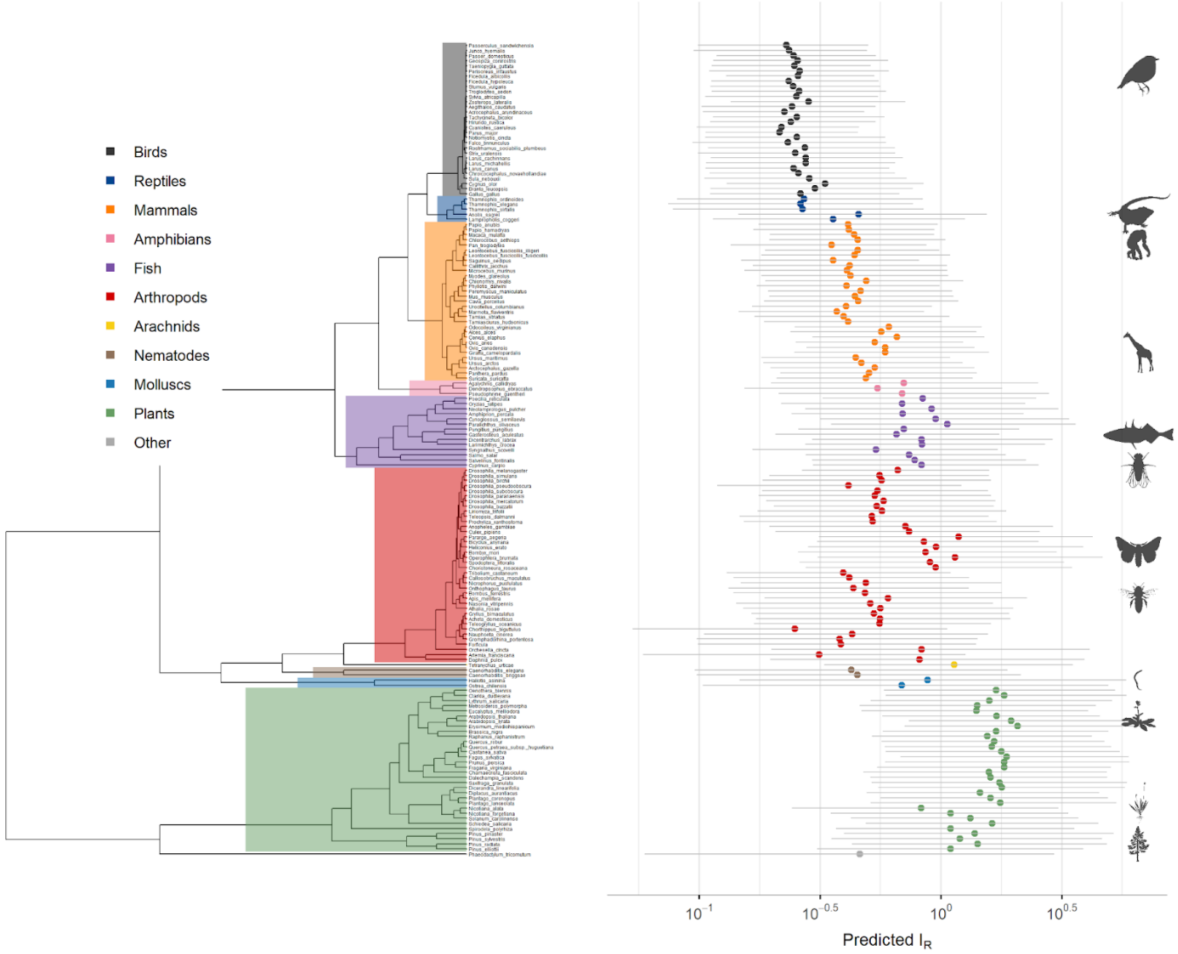


**Figure S10.** The predicted IR (mean scaled residual variance) from our model for a linear, morphological trait estimated with an animal model from a wild population for each species, plotted across the phylogeny with their 95% credible intervals.
